# Longitudinal and cross-sectional selection on flowering traits in a self-incompatible annual

**DOI:** 10.1101/2025.04.30.651553

**Authors:** Laura Leventhal, Karen J. Bai, Madeline A. E. Peters, Emily J. Austen, Arthur E. Weis, Jennifer L. Ison

## Abstract

Net selection on a trait reflects the association of phenotype to fitness, across an entire life cycle. This longitudinal estimate of selection can be viewed as the summation of selection episodes, each characterized by a cross-sectional estimate. Selection may be consistent in direction and strength across episodes for some traits, fluctuating in others, and for some, concentrated in a single intense event. Additionally, while selection on plant reproductive traits is predicted to be stronger through male fitness than female fitness, male fitness remains less studied. We investigated how selection on flowering traits in *Brassica rapa* varied temporally and spatially by measuring male reproductive fitness in four experimental populations with two spatial arrangements. To estimate longitudinal and cross-sectional selection, we introduced plants at successive intervals within a single reproductive season. We genotyped over 3000 plants and calculated selection on flowering time, duration, and total flowers. Cross-sectional analyses revealed directional selection was common, but patterns were masked by longitudinal estimates. Spatial population arrangement significantly impacted pollen movement, demonstrating how breeding timing and spatial aggregation interact to create complex evolutionary dynamics.

## Introduction

Selection is not a single stagnant parameter, and is instead a dynamic process that can fluctuate across many time scales. Within-season variation in selection is most effectively demonstrated by cross-sectional selection analyses, which measure selection on a subset of the population at a particular point in time (Endler, 1986; Kingsolver et al., 2001). While longitudinal selection studies reveal the net direction and intensity of selection by measuring the relationship between phenotype and lifetime fitness proxies, the cross-sectional approach can illuminate the mechanistic underpinnings of differential survival, mating, and reproduction throughout a breeding season, i.e., the *drivers* of longitudinal selection. Taken together, these approaches allow us to determine whether longitudinal selection arises through steady and consistent selection across an entire season, a few intense selection events at specific timepoints, or counterbalancing selection resulting in a longitudinal net-zero. Understanding how selection varies across shorter, episodic temporal scales can provide critical insights into how an ultimate, longitudinal association between trait and fitness came to be (Cockburn et al., 2008; Hoekstra et al., 2001; Losos et al., 2006; Walsh & Blows, 2009).

Unlike many animal species with highly synchronized breeding periods lasting only days or hours (Urquhart 1960; McDiarmid 1994; Schreiber and Burger 2001; Fogarty, Vollmer, and Levitan 2012), most flowering plants deploy reproductive structures sequentially over weeks or even months. Pollen transfer from donor to recipient is largely passive, mediated by biotic or abiotic vectors whose effectiveness fluctuates with environmental conditions (Ackerman 2000; Elzinga et al. 2007; Polatto, Chaud-Netto, and Alves-Junior 2014). Consequently, a plant’s reproductive success represents the sum of all day-to-day mating events within an entire breeding season weighted by their contribution to the mating pool. The association between trait values and realized mating (i.e., the direction and magnitude of selection on plant traits), could in theory vary across dates. Our experiment explores this within-season variation to determine the drivers of longitudinal selection. Variation in selection across dates could lead to several paths in which longitudinal selection is derived including 1) counterbalancing selection where two time points estimates counteract each other, 2) slow and steady selection, where most estimates within a season agree in direction and magnitude, and 3) a burst of selection where a one or a few strong events fully pull the longitudinal selection estimate in a single direction (**Fig. 1**). Many studies cannot evaluate reproductive success on such a fine scale due to time and effort constraints. To address this gap in the selection literature, we focus on the temporal scheduling of flower deployment and its impact on male reproductive success through pollen export over the entire breeding season. By dissecting the breeding season into discrete temporal windows, we can analyze how selection on male function varies day-to-day within a season, revealing how phenological traits shape reproductive outcomes across the flowering season.

**Figure 1.**
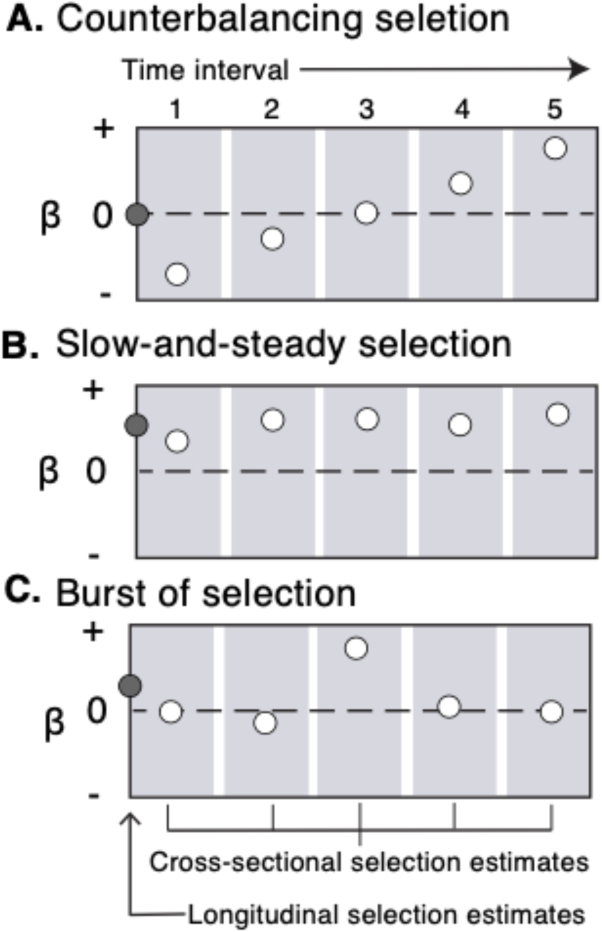
Conceptual diagram illustrating how longitudinal selection on traits can arise via several pathways of cross-sectional selection. In our theoretical population, selection on a given, unnamed, trait could theoretically arise through (A) counter-balancing selection where strong values in opposing directions counter each other and result in a net-zero longitudinal selection (B) slow-and-steady selection, where most selection estimates are of the same direction and similar magnitude, suggesting little variance in selection gradient estimates within the season, and (C) a strong burst of selection where overall longitudinal selection estimates are higher than the selection from the rest of the season would suggest, due to the sheer magnitude of the single interval’s selection estimate. Selection gradient estimates (β) overall for the season are the dark grey points while the cross-sectional estimates that comprise them are the white points within each time interval. The direction and magnitude of selection are important, where positive values indicate directional selection in the positive direction for that trait while negative values indicate directional selection in the negative direction for that trait, and points cross zero are non-significant and have no directional selection.

Natural fluctuations across a breeding season in population density of mates, competitors, predators, and mutualists could give rise to within-season variation in the direction or magnitude of cross-sectional selection, both hard (environmental selection) and soft (social selection) (Wallace 1975). Because these demographic shifts occur unevenly across space and time, the spatial aggregation of individuals within a population can influence ecological processes such as pollen movement and encounter distances, even if it does not directly shape selection pressures. For example, honeybees, *Apis melifera,* travel longer distances between flowers as spacing between individual plants increases, which can expand pollen dispersal and potentially increase outcrossing rates (Morris 1993). Additionally, when the population density of flowering individuals fluctuates, pollinators may be forced to forage over larger areas, further increasing mating distances. Temporal variation in factors like availability of mates can, in turn, influence selection patterns, as documented in animal systems (Kasumovic et al. 2008; Punzalan, Rodd, and Rowe 2010; Wacker et al. 2014; Gyulavári et al. 2017; Daupagne et al. 2023) and less frequently in plant systems (Parachnowitsch and Caruso 2008; Parachnowitsch et al. 2012). In flowering plants, density-dependent selection can be mediated by multiple, non-exclusive mechanisms. For instance, the density of competitors influences selection in flowering plants as competition for pollinators can arise when there is temporal overlap with blooms of competing species (Levin and Anderson 1970; Flanagan, Mitchell, and Karron 2011; Landry 2013; Richardson et al. 2021). Besides competitors, within-season variation in the pollinator community itself (composition, abundance, effectiveness) can influence selection (Fisogni et al. 2016; CaraDonna et al. 2017; Ison et al. 2018). Finally, a population’s flower density varies as plants enter and exit the flowering season and the individual-level floral display size rises and falls over time, which may affect both among- and within-individual competition for resources (e.g. Thomson 1980; Grindeland, Sletvold, and Ims 2005; Makino, Ohashi, and Sakai 2007; Feldman 2008; E. J. Austen, Forrest, and Weis 2015). If male fitness (seed siring success) is limited by access to mates, we might expect selection through male fitness to be particularly vulnerable to within-season changes in conspecific density and aggregation and pollinator service.

Despite the heightened sensitivity of male fitness to these ecological dynamics, our understanding of selection in plants remains disproportionately shaped by data on female fitness. This is largely because measuring male fitness– often dependent on elusive pollination and siring success– poses major practical hurdles, while measuring female fitness through seed production is more straightforward. This bias was illustrated in a meta-analysis of selection on flowering time, where only 5.7% (5/87) of studies evaluated selection via male fitness (Munguía-Rosas et al., 2011). Male and female fitness can differ in the degrees to which they are affected by resource availability, mating opportunity, and other factors (Bateman & Innes, 1948), and there may be a disagreement among selection estimates depending on the fitness input (Conner et al., 1996; Delph & Ashman, 2006; Hou et al., 2024; Tsuji et al., 2020). Notably, when male and female fitness optima for a trait differ, evolutionary outcomes in cosexual populations, such as in *B. rapa*, are expected to optimize total fitness (male plus female), possibly at the expense of strictly maximizing fitness for either sex alone. The shape of the male and female fitness functions could result in selection for a trait value range rather than a single optimum point.

One difficulty in estimating selection on the flowering schedule through the pollen-export fitness component arises when female qualities vary across time. Pollen export is intrinsically a trait of the donor, but must be quantified through the number of seeds sired. This poses no problem if all females are of equal and constant quality, i.e., invariant in the probability that pollen receipt results in offspring production. However, flowers produced toward the end of the flowering season typically have reduced fruit maturation rates (Weis and Kossler 2004; Austen, Forrest, and Weis 2015). This decline can stem from priority effects: early flowers access the full pool of maternal resources, which become progressively depleted for later blooming flowers. The ‘gene-trap’ method developed by (Devaux and Lande 2010) and refined by (Ison and Weis 2017) can help circumvent the priority effect bias. This approach involves introducing successive batches of pollen-recipient plants– all at identical developmental stages– into the experimental population, followed by paternity analysis of their seeds. By reducing variance among pollen recipients, this method reveals the strictly male component of selection through analysis of covariance between donor traits and siring success. Further, if the gene-traps are male sterile, they need not be considered sires of one another in analyses.

### Research goals

In this study, we investigate how selection on flowering schedule components – flowering time, flowering duration, and total flower production– via pollen export varies throughout the breeding season in *Brassica rapa* using the gene-trap method. Our primary research goal is to evaluate if season-long (longitudinal) selection gradients emerge from consistent selection across time, from episodic bursts of selection, or from counterbalancing directional changes in selection (all forms of cross-sectional selection). We predict that selection on phenological traits such as flowering time and duration will arise through counterbalancing selection across intervals, reflecting the changing phenotype status of mates throughout the season. We also test whether spatial aggregation of plants alters these patterns. Because pollinators travel greater distances when flowers are more widely spaced, we predict that populations with greater spacing between aggregates will exhibit longer pollen-dispersal distances, greater mate diversity, and potentially weaker spatial autocorrelation in siring success compared to closer-spaced populations. However, given the small spatial scale of our design, we expect the overall strength and direction of selection to be largely consistent between spatial treatments. This integrative approach provides insights into the basic evolutionary processes shaping flowering traits and contributes to our knowledge of how selection via pollen export operates in plant populations.

## Methods

### Study species: *Brassica rapa*

*Brassica rapa* is a non-clonal annual in the Brassicaceae. The species is native to Eurasia but is naturalized throughout much of North America (Gulden, Warwick, and Thomas 2008). Flowers are typically perfect (containing both pollen-producing and ovule-producing structures within the same flower), and plants from our seed source begin flowering 30-60 days after seed germination (Ison and Weis 2017; Austen and Weis 2016a). The inflorescence is an elongated raceme, and flowers will persist for one to three days but senesce more quickly once fertilized. Flowering can continue through the fruit set of the earliest flower (Austen and Weis 2016a). *Brassica rapa* is an obligate outcrosser with a sporophytic self-incompatibility system (Bateman 1955; Murase et al. 2020). Insect pollinators include generalist bees (Apidae, Halictidae), flies (Syrphidae), and butterflies (Lepidoptera).

### Experimental design

#### Field populations

We established four perfect-flowered experimental populations of *B. rapa* at the University of Toronto’s Koffler Scientific Reserve (44°1′N, 079°32′W) during summer 2011 using seeds bulk collected from a naturalized population in Eastern Quebec in 2009 (Austen & Weis, 2016a). We planted populations in one of two spatial aggregations (even or clumped, two populations per type), allowing us to determine the effect of mate density on selection and distance pollen traveled. Both spatial aggregations consisted of a hexagonal grid with 4 m sides. In the even aggregations, plants were evenly spaced with 0.5 m between nearest neighbors (211 total plants; **Fig. 2A**). The clumped arrangement had six small hexagons within a larger hexagon, with small hexagons 2.7 m apart and nearest neighbors within the small hexagons 0.17 m apart (216 total plants; **Fig. 2B**). Due to logistical and space constraints, the experiment was completed in two experimental runs over two time points. Populations in the first experimental run (even 1 and clumped 1) flowered July 4 – Aug 11. Populations in the second experimental run (even 2 and clumped 2) flowered Aug 16 – Oct 13 (even and clumped 2). Population plants (potential sires) for the first experimental run were directly seeded into the plots (two seeds per position and thinned when the first true leaves were present). Population plants for the second experimental run were planted in a three-season greenhouse and were transplanted into the plots right before the first flower (see Ison and Weis 2017 for more information). We monitored pollination visits to document changes in pollinator service across the experimental replicates. In total, our design included four experimental populations (i.e. even 1, clumped 1, even 2, clumped 2).

**Figure 2.**
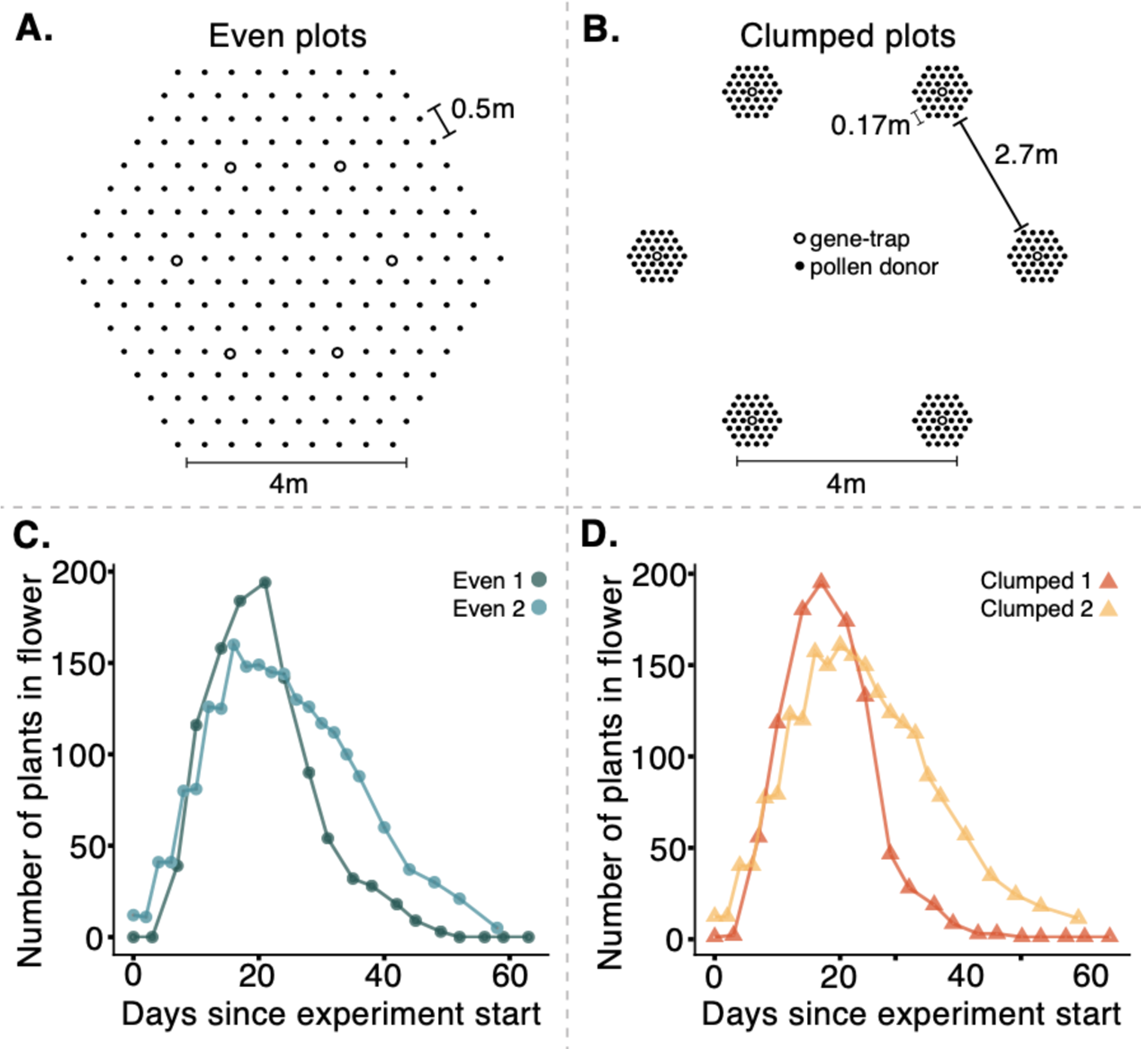
Diagrams of the experimental population spatial arrangements and flowering phenology. Populations were hexagonal 4 m sides. Each population had six gene-trap (male-sterile plants) positions denoted by an open circle **(AB)**. Gene-trap plants were added and removed from populations in consecutive batches every 3-5 days throughout the experiment (7 batches for even and clumped 1 experiment July-Aug time period, and 8 batches for the even and clumped 2 experiment Aug-Oct time period, see **Table S1** for placement schedule). For the even spatial arrangement **(A)** plants were spaced 0.5 m apart. In the clumped arrangement **(B)** there were six smaller hexagons at the corner of each of the population hexagons. Within each smaller hexagon plants were 0.17 m apart and smaller hexagons were 2.7 m from each other. Flower counts were recorded for all plants on census days corresponding to each data point shown in the flowering phenology curve **(CD)**. The even populations are denoted by circles and the clumped populations are denoted by triangles.

#### Gene-trap implementation

To assess male fitness and pollen movement, we deployed successive batches of male sterile plants (‘gene-traps’) as standardized pollen recipients. We produced these male-sterile plants by back-crossing Wisconsin FastPlants line 1-107 (carrying male sterility) into the Eastern Quebec population background (see Ison & Weis 2017: ‘constructing the gene-trap line’). This breeding approach yielded functionally female plants that were morphologically and developmentally similar to the perfect-flowered plants while maintaining male sterility. Male sterility of gene-traps is beneficial because male-sterile gene-traps do not need to be included as potential sires to one another in analyses. We planted cohorts of gene-traps plants in a 3 season greenhouse at the field station every week (see Ison & Weis 2017 for more information on growing conditions), and chose individuals who had most recently started flowering for deployment in the experiment.

Each population contained six gene-trap positions where male-sterile plants were placed. These positions were evenly spaced in the even populations and positioned at the center of each small hexagon in the clumped populations (**Fig. 2**). Gene-trap plants were systematically replaced approximately 3-5 days throughout the flowering season (**Fig. 2** and **Table S1**). The gene-traps that were placed in the populations were male-sterile plants that were early in their flower phenology (started flowering appropriately 2-4 days before they were placed into the populations). The temporal rotation of new male-sterile plants as pollen recipients at each interval eliminated variance in mate quality available to prospective sires across the season, ensuring our selection estimates reflected differences in pollinator service and/or in the competitive abilities of the perfect-flowered pollen plants as pollen donors. We genotyped the seeds produced by gene-traps to estimate siring success of the perfect-flowered pollen donors through paternity analysis.

We conducted five minute observation periods for each gene-trap position and four nearby perfect-flower plants during each gene-trap interval (30 plants observed per population per gene-trap interval, 15 h total observation). The number of visits observed did not differ notably by population, or by spatial aggregation within a replicate. However, we observed honeybees (*Apis mellifera*) and bumblebees (*Bombus spp.*) visiting experimental run 2 populations (**Fig. S1, Table S3**), indicating that the pollinator assemblage shifted somewhat across the two time periods. We did not record the time each pollinator spent on the flower.

### Phenotypic measurements and genetic analysis

We recorded three flowering traits for each pollen-donor (854 perfect-flowered plants): day of first flowering (flowering time), flowering duration, and the total number of fresh flowers (summed across census days). Prior to analysis, we standardized all phenotypes by z-scoring them within each of the four populations.

We collected tissue samples from all perfect-flowered plants (potential sires) and each gene-trap plant, and dried in silica gel. During winter 2011-2012, we germinated and grew gene-trap offspring in a greenhouse (see Ison and Weis 2017 for methods). For this study, we aimed for twelve offspring per gene-trap plant but some gene-trap plants produced fewer than twelve viable seeds and for a few gene-trap plants, we sampled more than twelve offspring (mean 10.9; 1970 total offspring; **Table S1**). We took tissue samples from the offspring when at least two true leaves were present, and dried them in silica gel.

#### Genotyping methods

We extracted DNA from the dried tissue samples using isopropanol precipitation methods (Rogers, Burns, and Parkes 1996). We characterized the plants’ DNA at ten *B. rapa* microsatellite loci (Suwabe et al. 2002; Lowe et al. 2004) (for primer information, see **Table S2**). We conducted 10 ul polymerase chain reactions with 1 ul of DNA template, 5 ul of GoTaq Colorless MasterMix (Promega), 0.5 ul of BSA, 0.25 ul of additional MgCl_2_, 0.16 ul M13-tagged fluorescent dye (ABI dye set-33), 0.05 ul of forward primer with M13 tail, 0.6 ul reverse primer, and DNase free water to volume. Each PCR consisted of a 3 min 94°C denaturing cycle followed by 30-32 cycles of 30 s at 94°C, 30 s at annealing temperature (see **Table S2**), and a 30 s extension step at 72°C. Three additional cycles were run with an annealing temperature of 55°C followed by a final 10 min 72°C extension step. We sent the PCR products to the University of Toronto’s Center for Analysis of Genome Evolution and Function for capillary gel fragment analysis. We scored the resulting genotypes using Peak Scanner version 1.0 (Biosystems 2006). To calculate genotyping error, we repeated the PCR and fragment analysis on 96 haphazardly selected samples. We calculated both the allelic dropout error rate, ε1, and the stochastic error rates due to typing error, ε2, which can be attributed to mutation, miscalling, polymerase error, and data entry error (Wang 2004).

### Paternity Analysis/ MCMC Model

We conducted analyses using R 4.4.0 (R Development Core Team 2024). We assigned proportional paternity, reconstructed pedigrees, and checked for convergence using MasterBayes and coda for analysis (Hadfield 2017; Plummer 2024). MasterBayes uses Bayesian statistics to find the maximum likelihood of paternity through the calculation of a posterior distribution relying on both genetic and phenotypic data to find the highest probability offspring-paternal pair (Jones et al. 2010; Hadfield 2017). The statistical inference employs a Bayesian probabilistic architecture to simultaneously derive a posterior probability distributions across 1) genealogical relationships represented in the pedigree structure *P, and* 2) the coefficient vector β that parametrizes the log-linear model incorporating non-genetic covariates. This joint estimation process integrates the uncertainty in both the familial relationships and the magnitude of non-genetic effects within a unified probabilistic framework. The non-genetic information that is an input of the model comes out as the multinomial log-linear model. The coefficients from the multinomial log-linear model are analogous to the coefficient of a multiple regression, thus the coefficient derived on non-genetic data from paternity assignment of offspring outputs can be interpreted as a selection gradient (see Austen & Weis, 2016a).

This Bayesian approach to paternity assignment allows for us to retain more offspring than with a frequentist paternity assignment approach because the frequentist approach removes offspring whose potential sires do not pass a confidence threshold. Losing this valuable data can reduce the sample size and potentially bias selection estimates. More critically, the frequentist method treats the paternity assignment as an error-free measurement which can propagate uncertainty and violates the principle of single-stage inference (Hadfield et al. 2006). In contrast, the Bayesian approach simultaneously estimates paternity probabilities and selection gradients which allows us to retain all of the offspring and accounts for uncertainty when estimating selection values. Further, this approach allows the selection gradient estimation to influence the paternity estimates and vice-versa, which thereby acknowledges that there can be error on both parts (paternity and selection gradients) and simultaneously optimizes both to maximize the likelihood of the observed offspring. Both the frequentist and Bayesian approach to paternity assignment are valid and useful, but for the scope of our study, we found the Bayesian approach to be most appropriate.

To parameterize our model, we initialized three independent MCMC chains with differing starting conditions, which is standard practice for Bayesian inference to ensure through exploration of parameter space and to avoid local optima. The first two chains were initialized with fixed genotyping error rates (ε1 and ε2) and allele frequencies, while the third chain incorporated normally distributed random starting values for the beta coefficients, with a mean vector of [-0.1,0.1] and a standard deviation of 0.05. We optimized MCMC sampling using a tuning parameter of 0.325 to achieve acceptance rates between 20% and 50%. Each chain was run for 110,000 iterations with a burn-in period of 10,000 iterations. To reduce autocorrelation, we employed a thinning interval of 100, resulting in 1,000 store iterations per chain. The chains were run with a maximum mismatch tolerance of 5. This multi-chain approach with different initialization parameters allowed us to assess convergence and mixing of the MCMC sampling while accounting for potential genotyping errors in the pedigree reconstruction. We tested all chains within a model for convergence using the Gelman-Rubin Diagnostic using the coda package (M. Plummer et al. 2006), and removed all models from the analysis that did not reach convergence (convergence is indicated by values lying between 0.9-1.1). In total we had 46 unique models (4 multivariate longitudinal, 30 multivariate cross-sectional, and 12 univariate longitudinal [to evaluate total selection, see **Fig. S2**]) with three chains each.

### Selection analysis framework

#### Measuring longitudinal and cross-sectional selection

We estimate selection gradients (β) on our phenotypes for each interval (cross-section) and for the entire season (longitudinal). We calculated longitudinal selection gradients by randomly subsetting the offspring produced in a given gene-trap interval by the proportion of plants in flower and then using the model to measure overall selection for the season. We did this random subsetting to account for temporal variation in pollen availability. For the cross-sectional approach, we estimated selection in discrete intervals across the flowering season but we restricted the pollen pool to specific windows when a single batch of gene-trap plants were present in the field (7 intervals for even and clumped 1 populations and 8 intervals for even and clumped 2 populations; **Table S1**). Here, potential sires were limited to plants producing pollen during the active gene-trap interval and thus plants that were non-flowering were removed from the interval-specific analysis. This approach generated interval-specific selection gradients that revealed how selection pressures shifted throughout the flowering season; note that traits were standardized at the population level (not within temporal windows).

Our multivariate selection framework incorporates all three flowering traits (flowering time, flowering duration, and total flower number) simultaneously into the log-linear model, providing an estimate of direct selection on the focal trait. This multivariate approach disentangles the network of trait associations, allowing us to isolate the direct selection acting on each trait while statistically controlling for the influence of correlated traits. Thus, the multivariate selection model partitions the shared variance of traits to reveal which traits are directly under selection versus those that experience indirect selection through correlation with other traits. We additionally measured indirect selection on each trait using univariate models where only the trait of interest was incorporated into the log-linear model, providing a selection differential and a measure of total selection for both our longitudinal and cross-sectional analyses in order to compare direct and indirect selection which may create conflicts due to trait correlations (**Fig. S2**).

### Distance of pollen movement analysis

To determine how far pollen was moving in the two population spatial aggregations and at different points in the season, we used a log-linear model framework to estimate the spatial effect coefficient. The log-linear model framework allows for the incorporation of spatial effects through pairwise distances between maternal plants and potential sires. The spatial effect coefficient (β) from the posterior distribution quantifies the decay in mating probability with increasing distance and follows an exponential decay function (Exponential decay = e^β*x^, where β is the peak of the posterior distribution and *x* is the pairwise distance). The spatial component was incorporated directly into the multinomial log-linear model with MasterBayes. The exponential decay model provides a mechanistic understanding of pollen movement where negative β values indicate decreasing probability of successful mating with increasing distance between pairs of plants. We measured the distance decay coefficient for each population overall in the season and also at each time interval within each population.

## Results

We successfully genotyped nearly all pollen donors (n=854), gene-trap (n=264), and offspring (n=1970) plants at the 10 microsatellite loci. There was 3.7%, 3.8%, and 2.5% of the genotypic data missing across loci, for pollen donor, gene-trap, and offspring plants, respectively. For the Bayesian approach to paternity assignment, we ran three chains with different parameters per model to ensure proper mixing. We tested convergence of our three chains per analysis and found that 97.8% (45/46) of our models converged across the three chains. Analyses that did not reach convergence with the Gelman-Rubin Diagnostic were removed from the results.

Within each population, there was variation in the number of successful dads for which we genotyped the offspring. Even 1 and even 2 had 100 and 114 paternal plants represented respectively, and clumped 1 and clumped 2 had 96 and 117 paternal plants represented respectively. The average number of offspring per dad remained relatively constant across populations and across populations and averaged to 1.67 offspring per sire (**Table S4).** Such low values were not unexpected given the total population size and our sampling scheme. Across all populations for the entire season, maternal gene-trap plants had a range of 0 to 15 unique sires as potential mates (**Table S5**). Even 1 and even 2 populations had an average of 3.71 and 3.94 unique sires per gene-trap, and clumped 1 and clumped 2 had an average of 3.36 and 3.79 unique sires per gene-trap.

### Longitudinal selection on flowering traits via pollen export

Selection varied among populations and traits when measured across the entire flowering season. One population (even 2) showed significant directional selection favoring late flowering (β [95% CI] = 0.040 [0.009,0.072]), while the remaining populations did not show significant evidence of selection on flowering time (**Fig. 3A**). We observed significant longitudinal selection for longer flowering duration in one population (clumped 2, β [95% CI] = 0.037 [0.020,0.055]) but no significant selection in the other populations (**Fig. 3B**). Selection favored more total flowers in three populations, with only the clumped 1 population having no significant selection (β [95% CI] = 0.007 [-0.004,0.017], **Fig. 3C**).

**Figure 3.**
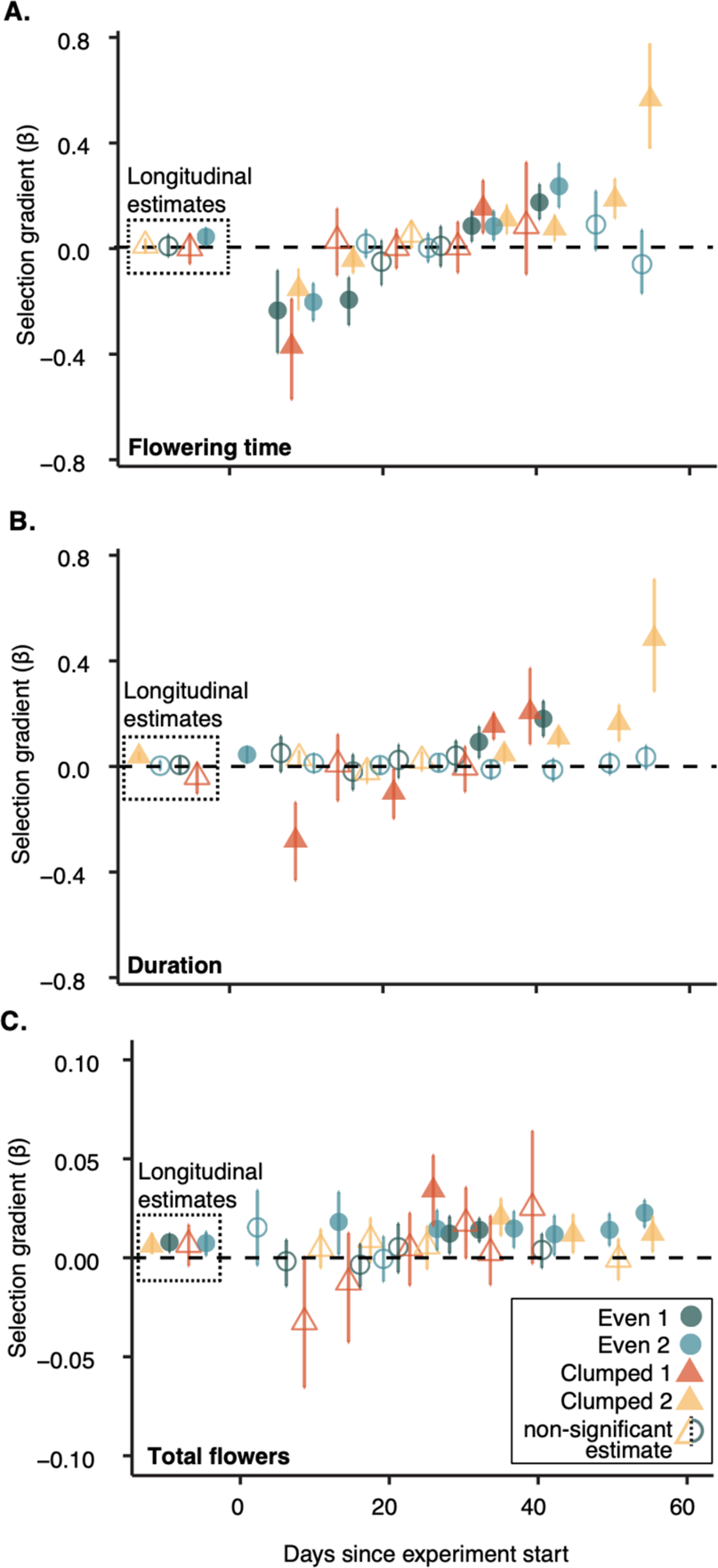
Longitudinal and cross-sectional estimates for selection gradients (95% CI) on **(A)** flowering time, **(B)** duration of flowering, and **(C)** total flowers in four experimental populations. Longitudinal selection gradients are depicted in the dashed box, while selection gradients in each time interval correspond with the days since being placed in the field axis. Blue circles correspond to evenly spaced populations and yellow and orange triangles represent clumped populations. A filled-in shape indicates the estimate and 95% CI do not cross zero while an empty shape indicates the estimate and 95% CI cross zero and are therefore non-significant estimates.

### Cross-sectional selection on flowering traits via pollen export

Cross-sectional selection analysis using the gene-trap sampling periods revealed dynamic patterns of selection that varied both temporally and spatially. All populations showed temporal shifts in selection on flowering time, generally favoring early flowering at the beginning of the season and late flowering towards the end, with non-significant estimates during mid-season (**Fig 3A**). Cross-sectional selection gradients on flowering time had the largest magnitudes in clumped populations at the beginning and end of the flowering period **Fig. 3A, Table S6**).

Cross-sectional selection on flowering duration and total flowers varied based on the population spatial aggregation. Both spatial aggregations had modest selection for longer flowering duration in many of the gene-trap intervals with stronger selection in end-of-season plants (clumped 2, (β [95% CI] = 0.484 [0.285,0.709], **Fig. 3B**). The clumped 1 population was the only population that showed selection for shorter duration (β [95% CI] = -0.280 [-0.431, -0.137] and β [95% CI] = -0.100 [-0.197,-0.009]). For total flowers, the even populations had relatively consistent selection for more total flowers throughout the season (**Fig. 3C, Table S6**), and there was never a significant cross-sectional estimate for fewer total flowers.

### Pollen movement in differing spatial arrangements

Pollen movement patterns varied by spatial arrangement across the flowering season. Clumped populations generally showed a smaller distance decay coefficient than even populations throughout the season (even 1 β [95% CI] = -1.19 [-1.38,-1.00], even 2 β [95% CI] = -0.59 [-0.72,-0.47], clumped 1 β [95% CI] = -0.30 [-0.38,-0.23], and clumped 2 β [95% CI] = -0.42 [-0.50,-0.35], suggesting that pollen moves farther in clumped populations (**Fig. 4A, Table S7**). The even populations had more variation in their decay coefficients across the season, and distance decay was largest during peak flowering (pollen moved the shortest distances), while clumped populations had a relatively consistent distance decay coefficient throughout the season (**Fig. 4B, Table S7**). Because we calculated the exponential distance decay with a Bayesian model, the distances predicted are an extrapolation and may not be plausible distances given the experimental set up (i.e., a distance combination might not exist, but will be predicted by our model). Further, the maximum distance pollen can move in an even population is 6 m while the maximum distance is 7.5 m in a clumped population.

**Figure 4.**
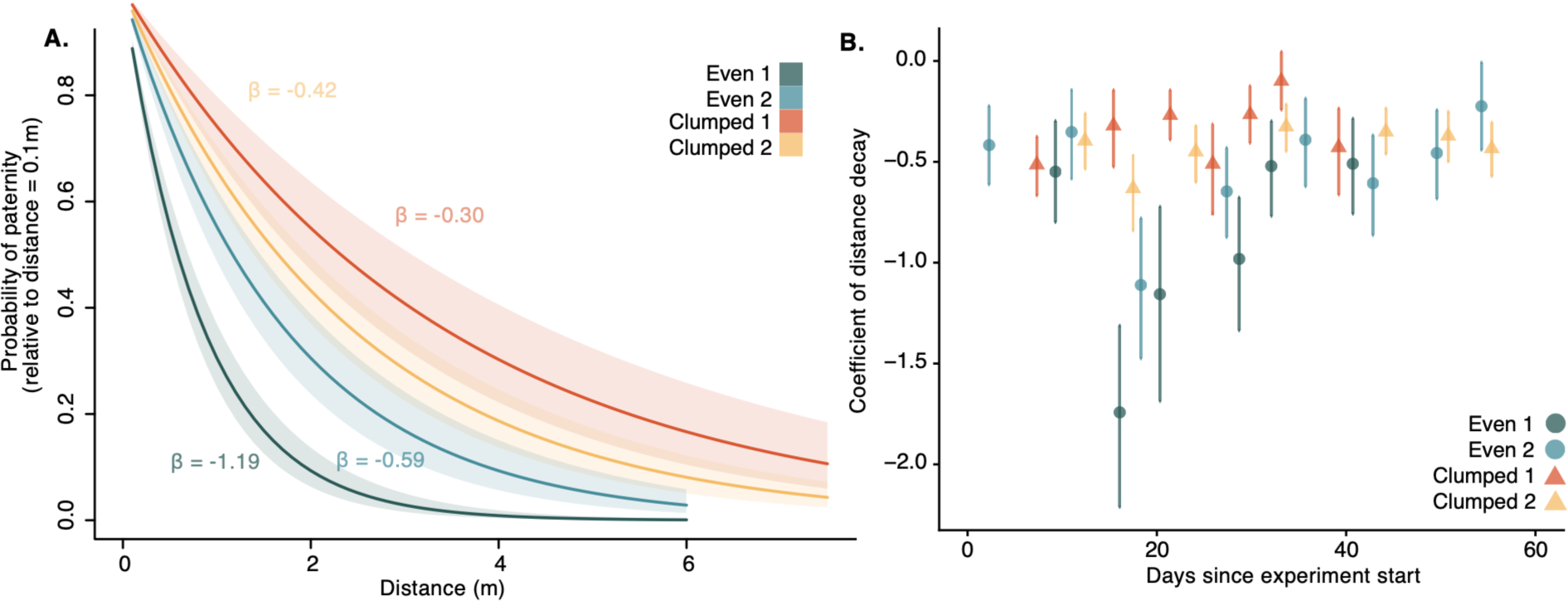
**(A)** Probability of paternity relative to distance (m) for the two population types plotted as an exponential decay of the distance decay coefficient. Shaded regions represent 95% Bayesian Confidence Intervals. Blue colors represent even populations and yellow and orange colors represent clumped populations. Clumped populations cut off a max distance at 7.5 m while even populations cut off max distance at 6 m because that is the greatest distance pollen could travel from sire to gene-trap in the respective spatial aggregations. **(B)** Coefficient of distance decays over cross-sections of time and plotted without exponential decay function, error bars represent 95% CI. In both panels, the blue shades represent even populations 1 and 2, while the orange and yellow shades represent clumped populations 1 and 2.

## Discussion

### Cross-sectional selection analyses reveal mechanisms of longitudinal selection in a plant with a long-breeding season

Longitudinal analyses capture season-wide trends, but fine-scale temporal analyses reveal shifts in both strength and direction of selection, clarifying how season-wide gradients emerge (**Fig 3**). We saw similar selection gradient trends over time for our four populations regardless of the spatial aggregation, suggesting that the selection was not strongly affected by differences in fine-scale density of conspecifics. First, we aimed to determine how selection on our three focal traits occurs by dissecting the longitudinal selection estimates with weighted cross-sectional estimates. In principle, seasonal selection estimates could arise from several paths including slow-and-steady selection, a burst of selection at one or a few time intervals in a season, or counterbalancing selection,where differing directions of selection at similar magnitude negate the longitudinal effect (**Fig. 1**).

Cross-sectionally early season intervals favor early flowering and late season intervals favor late flowering. This suggests that longitudinal selection on this phenological trait arises through counterbalancing selection (**Fig. 3A**). Interestingly, however, in one of the populations we found longitudinal selection for late flowering, indicating that counterbalancing patterns are not universal across sites. Cross-sectional estimates following phenology of the population may seem expected because the gene pool at the beginning of the season is dominated by early flowering individuals. Other non-exclusive possibilities include a true advantage of being among the very first pollen-producing plants, or demographic effects of being a part of the mating pool for the entire interval. Selection for the earliest flowering plants could arise due to reduced competition for pollinators, potentially increasing reproductive success (Elzinga et al., 2007; Kehrberger & Holzschuh, 2019). In mid- and late-season intervals, individuals remain in the gene pool from earlier cohorts. We found stronger selection for late flowering at the end of the season, which could suggest either a decline in the quality of pollen from early- and mid-flowering plants (Ashman & Schoen, 1994) or a genetic advantage of late-flowering individuals across contexts (Austen & Weis, 2016a). Experimental tests of pollen quality across flowering time cohorts would help resolve this mechanism. Selection on flowering time may also be influenced by ecological factors other than mate availability and quality such as resource availability (Forrest and Thomson 2010), competition (Mosquin 1971), and herbivory (Pilson 2000).

Flowering time has a high heritability (e.g. Dorn & Mitchell-Olds, 1991; Fox, 1990; Geber & Griffen, 2003; Reed et al., 2022), and with our Quebec populations of *B. rapa* previous studies have estimated narrow-sense heritability of flowering time at 51%, 57%, and 61% (Austen & Weis, 2016a; Ison & Weis, 2017). The strong heritability of flowering time could make longitudinal estimates of selection highly informative about potential evolutionary responses in these populations. Cross-sectional selection estimates, rather than representing competing processes, are best understood as components of temporal segments that together make up the longitudinal estimate. The decomposition enables finer-scale resolution of how selection operates through different intervals or ecological contexts.

Although flowering duration is often correlated with heritable traits such as flowering time, its own heritability remains poorly characterized due to limited direct measurements in selection studies. Existing studies report moderate to low heritability for flowering duration (Bhakta et al., 2017; Cheng et al., 2006; Hof et al., 1999; Williams et al., 2022). Theoretically, extended flowering duration can enhance male fitness by increasing mating opportunities for pollen transfer, yet empirical evidence is scarce (Austen & Weis, 2016a; Lloyd, 1980; Xu, 2024). However, longer duration could incur resource costs that create trade-offs generally such as longer duration resulting in lower quality flowers (Ashman & Schoen, 1994) or antagonism between male and female fitness components (Ashman & Morgan, 2004; Zhao et al., 2020). In our experiment, the relationship between duration and fitness varied, but generally selection favored longer duration later in the flowering season (**Fig. 3B**). Specifically, selection gradients had larger magnitudes late in the flowering season than early and mid-season, indicating counterbalancing selection resulting in nonsignificant longitudinal estimates, except in one population (clumped 2). There were even instances of selection for a shorter duration early in the season in clumped 1, while later in the season for that population there was selection for longer duration, highlighting the impact of counterbalancing selection gradients.

In contrast with duration and flowering time, which have longitudinal selection arising through counterbalancing cross-sectional selection, selection on total flowers was characterized by having consistent, steady, and small selection for more total flowers (**Fig. 3C**). The magnitude of selection gradients for total flowers is significantly smaller than on flowering time or duration (**Fig. 3**).Our analysis measured selection on total flowers independent of the effects of duration (because we used a multivariate model, accounting for trait correlations), thus total flowers here could be a proxy for display size. Theory predicts an intermediate optimum for display size because too large or attractive of a display size may result in wasteful geitonogamous transfer (i.e., pollen transfer occurring from different flowers on the same individual) (Robertson and MacNair 1995), which may indicate why our estimates are low. A low estimate for a potentially morphological trait contrasts to a finding from a selection meta-analysis where researchers found that selection on morphological traits had larger selection estimates than phenological traits (Kingsolver et al. 2001). Similar to flowering duration, heritability estimates of total flowers are varied and much less common than estimates for flowering time (Drennan et al., 1986; Fogaça et al., 2012; Hof et al., 1999) and thus the evolutionary implications of selection for more total flowers in this species remains unknown.

Although our analyses provide insights into trait-specific selection, we recognize that part of the observed variation in flowering traits could reflect underlying differences in overall plant condition. Previous studies in *B. rapa* have shown that measures of plant size, such as height and stem diameter, are often strongly correlated with flower number and total reproductive output (Austen et al. 2015). Large, more vigorous individuals generally produce more flowers, potentially contributing to greater total flower counts and extended flowering durations, although these attributes can have tradeoffs for male fitness as we discuss below. Due to logistical constraints of adding more measurements, we were not able to include direct measures of plant size as covariates in this study. Future work incorporating direct measures of size and physiological status would help to test how much observed selection on flowering traits reflects differences in conditions versus intrinsic variation in phenological or developmental timing.

### Insights into male-mediated selection on floral traits and phenology

Surprisingly, we found small magnitudes of selection for more total flowers in all populations across time points (**Fig. 3C**), which may be linked to potential costs to male fitness in having more flowers. While theoretical models suggest that producing more flowers should increase opportunities for pollen dispersal and thus male reproductive success, plants often face resource allocation constraints that force trade-offs between flower quantity and quality (Lanuza et al. 2023, Burd 1999) Further in self-compatible species (unlike *B. rapa* which is self-*incompatible*), there may be a tradeoff with total flowers and male fitness because of geitonogamy, where a flower is pollinated by pollen from a different flower on the same individual (Robertson and MacNair 1995). If a plant has too many highly attractive flowers, there may be a cost to siring success, and therefore an intermediate display size or attractiveness may optimize both pollination visits and departings. These dynamics underscore the need to consider both resource-based trade-offs and phenological variation in selection studies while considering the specific mating biology of the species.

Much literature of selection on traits in flowering plants has focused on the mating biology of the female component, and less attention has been paid to selection via the male component (Munguía-Rosas et al., 2011). For flowering time specifically, most studies have evaluated selection via female fitness and found overwhelming selection for early flowering (reviewed in Austen et al., 2017; Austen & Weis, 2016b, Munguía-Rosas et al., 2011). Our study, which measured selection via pollen export on flowering time, found that in one population, longitudinally there was selection for late flowering (**Fig. 3A**)–a rarity in flowering time literature. Adding the cross-sectional selection perspective to flowering time in our study highlights that selection via pollen export sometimes favors late flowering in all four populations (**Fig. 3A**). Overall, the alternating advantages of early and late flowering in this experimental design, combined with evidence that female function likely favors early flowering (although cross-sectional selection on flowering time via female function has to date not been evaluated empirically), suggest a potential mechanism for minimal net directional selection on flowering time once the fitness components are integrated. While total fitness integrates both components of male and female fitness, our results demonstrate that male function via pollen export alone does not reinforce early flowering, challenging assumptions based solely on female function.

Researchers evaluating sexual selection conflict in a gynodioecious species, *Cyananthus delavayi*, demonstrated that when protandrous pollen was removed from perfect flowered individuals, the fitness of manipulated individuals was higher than if pollen was not removed and was equal to female-only individuals, highlighting that sexually antagonistic selection may play out differently depending on the breeding system and ecological context (Wang et al., 2021). Regrettably, female fitness (seed production) was unmeasured in our non-gene-trap plants, limiting our ability to directly test the male-female counterbalance hypothesis. In a co-sexual species, theoretically, different constraints to male and female fitness could create weakened selection through total (reproductive) fitness (Anthes et al., 2010; Ashman & Morgan, 2004; Bateman & Innes, 1948; Burd & Callahan, 2000; Campbell, 1989. However, even studies measuring both male and female success remain incomplete without accounting for unmeasured components such as correlations between flowering time genotype and seed variability or early-life mortality (Wadgymar et al., 2017; Wadgymar et al., 2024).

Our analysis acknowledges that male fitness is shaped not only by mating opportunity but also by intrinsic features of the pollen donor. This design seeks to reduce (though not fully eliminate) confounding effects of mating opportunity, allowing sharper inference on intrinsic determinants such as pollen quality. For instance, in *Medicago sativa* (alfalfa), pollen (Brunet et al. 2019) declines overall the plant’s flowering (Brunet et al. 2019), even shortly after the pollen is presented (Brunet et al. 2019). Additional selective components include the competitive ability of individual pollen grains to fertilize ovules post-deposition, reflecting a selective layer on pollen quantity and quality relative to competing grains from multiple sires (Erbar 2003). Pollen competition involves diverse traits such as optimizing pollinator visitation duration to avoid pollen packing, producing sufficient pollen volume to outcome rivals, and floral morphologies enhancing deposition on pollinator body parts most likely to contact stigmas (Delph 2019). Collectively, these intrinsic factors and competitive processes underscore that male reproductive success is impacted by more than mating frequency. Ultimately, a full understanding of trait evolution requires integrating male and female reproductive fitness together with often unmeasured fitness components, and we encourage future studies to consider this comprehensive approach.

### Pollen moves farther in clumped aggregations, especially early and late in the season

We observed striking differences in successful pollen movement between even and clumped spatial aggregations, where overall pollen moved farther in clumped than even populations (**Fig. 4A**). Although the maximum distances were different, if we compare the minimum largest distance (6m), we can see that pollen is still moving farther in clumped populations, which is consistent with other studies that found pollinators move farther when plants are farther apart (Morris 1993). While our design kept the overall number of plants constant between even and clumped populations, the aggregation of the plants and density immediately around the gene-traps varied (50 cm in even populations and 17 cm in clumped populations). Other studies have found that pollinator foraging behavior changes with plant density (Cresswell, 2000; Grindeland et al., 2005; Kunin, 1997). However, how these changes in foraging behavior impacts pollen movement distance is less understood. In one study, bumblebees were more likely to skip over plants and their foraging had less directionality when plant density increased but that did not lead to changes in pollen movement distances, based on foraging observations (Cresswell, 1997). In another study more analogous to the set up of our own, bumblebees had longer visitation rates in sparser (clumped) populations and preferred larger display sizes, which also drove the pollinators to travel farther distances potentially resulting from increased visibility (Mustajärvi et al. 2001). As pollinators of *B. rapa* receive nectar and pollen as rewards, it is worth noting that our gene-trap plants do not produce pollen. This may have affected the frequency or duration of pollinator visits. However, because our study focused on tracking pollen export from fertile plants, any reduction in visitation to gene-traps is unlikely to bias our measures of pollen movement and siring distances.

The effect of pollination distances being dependent on spatial aggregation is pronounced in the intervals of time too. The distance pollen moved in even populations varied across the season such that pollen moved the greatest distance early and late in the season and shorter distances at the peak flowering middle of the season (**Fig. 4B**). Such differences could be attributed to increased mating opportunities at peak. This is consistent with other studies that also found longer-distance pollen movements early and late in the season (Ison et al., 2014; Kitamoto et al., 2006). In contrast, pollen moved generally farther in clumped populations at most points in time, despite closer plant proximity, and clumped populations maintained consistent movement distances across the season (**Fig. 4B**). We might expect changes in selection estimates based on spatial aggregation due to pollinator preference and decisions clearly being invoked in distance traveled; however, selection estimates are consistent regardless of spatial density, pointing to the robustness and importance of these flowering traits.

### Conclusions and next steps

Our study demonstrates how multiple approaches for estimating selection can reveal complex evolutionary dynamics that might otherwise remain hidden. The cross-sectional approach reveals nuanced patterns of selection throughout the flowering season that were masked in longitudinal estimates, highlighting the importance of temporal scales in selection studies and displaying the mechanisms behind longitudinal selection estimates. By examining selection through male fitness in *B. rapa*, we uncovered selection for late flowering in one of four populations, which is a rare occurrence in the flowering time-selection literature. We found that total flowers had much weaker selection gradients but consistent selection throughout the season pointing to the mechanism of steady and consistent selection for more total flowers. Finally, spatial aggregation emerged as a key factor influencing pollen movement patterns, but did not impact selecting gradients, suggesting the robustness of selection on these traits.

Future studies should integrate both male and female fitness components to provide a comprehensive view of selection acting on plants. Additionally, incorporating temporal sampling designs will better capture how selection varies throughout the season, allowing for deeper insight into the drivers of selection. This multifaceted approach to studying selection will enhance our understanding of how natural populations respond to selection pressures across different spatial and temporal scales.

## Data Accessibility Statement

*The data underlying this article are available in: (doi for dryad if manuscript is accepted).* If the reviewers would like to see the code, please have the editor reach out to the corresponding author.

## Author Contributions

JLI and AEW conceived and designed the experimental populations and gene-trap method with input from KJB and EJA. JLI and KJB collected field and genotype data. LL led the analysis with help from EJA, JLI, MAEP and AEW. LL wrote the initial draft of the manuscript with contributions from JLI and AEW. All authors contributed to manuscript revisions.

## Acknowledgements

The authors thank BA, DC, JEI, JJ, GRS, YS, and GT for field and greenhouse assistance. JLI also thanks Raven for companionship in the field. CG, GRS, YS, GT, and JT, helped extensively with lab work and we are grateful for their efforts. We thank SMW for feedback on research design and ML, JP, KSM, RE, TB, ARP, and EVS, for feedback on early drafts of the manuscript. Initial funding was provided by grants from the Canada Foundation for Innovation, the Natural Sciences and Engineering Research Council of Canada (RGPIN-2016-06540), and the University of Toronto. Additional funding was provided through the College of Wooster’s Copeland, Luce, and Wilson Awards. While working on the manuscript JLI was supported by the College of Wooster’s research leave program and LL was supported by the National Science Foundation Graduate Research Fellowship (Fellow ID 2020304890).

## Supplemental Figures

**Figure S1.**
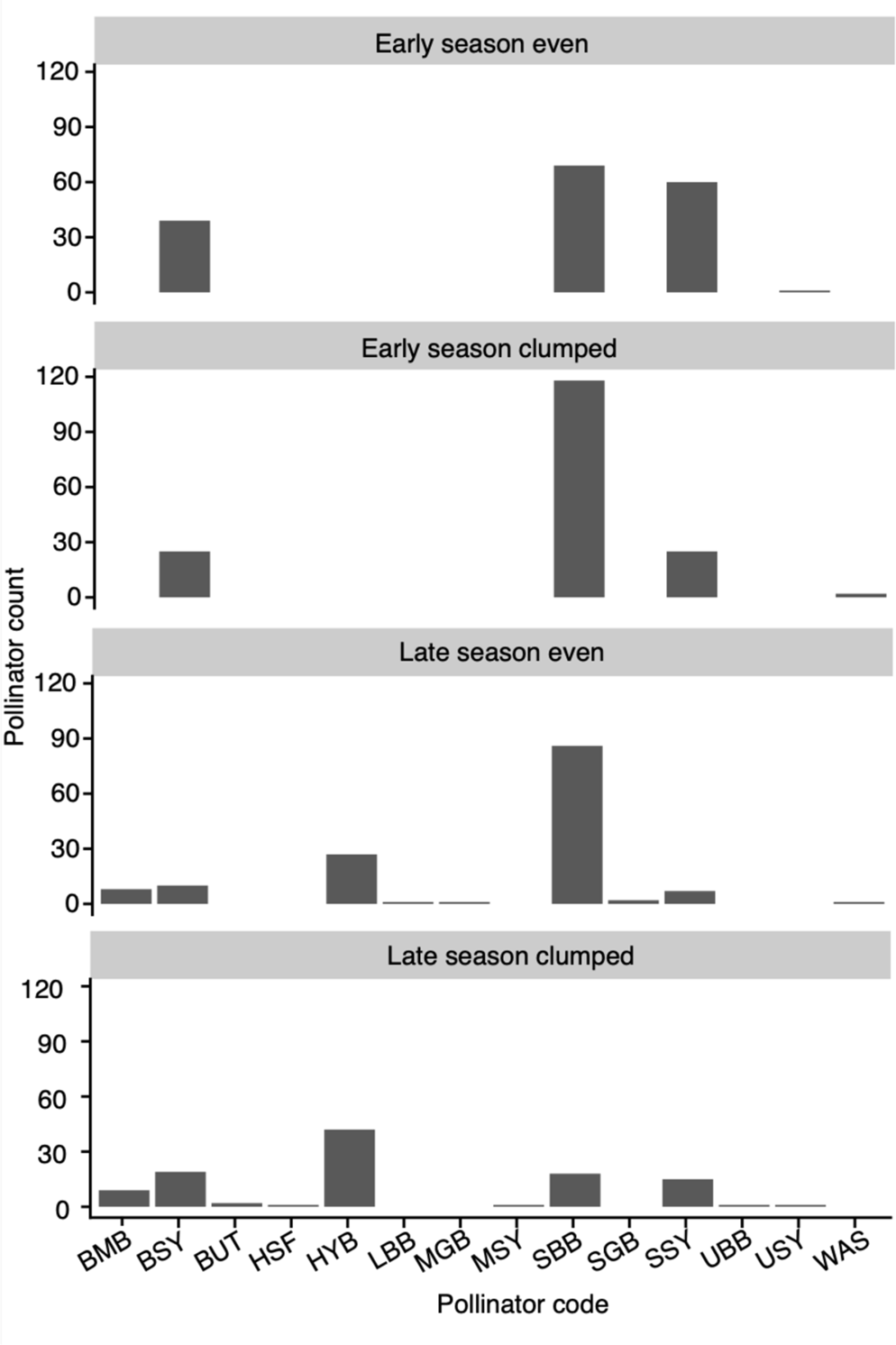
Pollinator counts by pollinator category for each of the four plots (BMB = bumblebee, BSY = big syrphid fly, BUT = butterfly, HSF = house fly, HYB = honey bee, LBB = large black bee, MGB = medium green bee, MSY = medium syrphid fly, SBB = small black bee, SGB = small green bee, SSY = small syrphid fly, UBB = unknown black bee, USY = unknown syrphid fly, WAS = wasp). Pollinator counts were obtained when we conducted six five-minute pollinator observation periods every four to eight days at each of the four plots. Each observation period included a gene-trap plant and the six closest plot plant positions. We recorded all insects that visited a reproductive structure on one of the observed plants. We did not collect the pollinators, so as to not impact the pollinator community.

**Figure S2.**
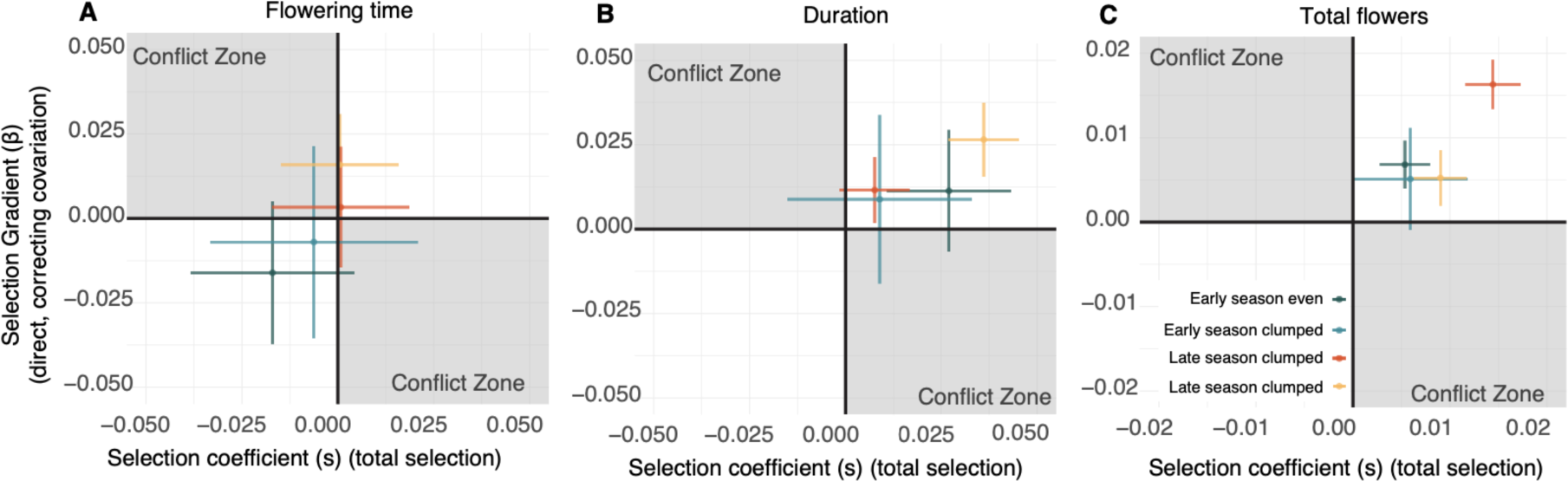
Overall selection conflict for three focal traits, lines represent 95% CI. Direct selection, which accounts for trait variation through a multivariate model is on the y axis, while total selection, which does not account for trait correlations and uses just a univariate model, is on the x axis. If total and direct selection are different signs (positive and negative) and significant (do not cross zero), then the trait in that population could be considered to be in a ‘conflict zone’ where selection on a trait could be mediated by correlations with other traits or suites of traits. We do not find any selection of traits in our populations significantly in a conflict zone. For an example of this analysis with true conflict zones see (Ruffley et al. 2023).

## Supplemental Tables

**Table S1.** Information about the gene-trap plant batches for early and late season populations. Batches are gene-trap plants that were placed into the experimental plots on the first day of the ‘gene-trap time period’ and removed on the last day. Approximately twelve offspring were genotyped from each of the six gene-trap plants from each batch.

| Gene-trap interval | Gene-trap time period | Gene-trap offspring genotyped (even plot) | Gene-trap offspring genotyped (clumped plot) |
| --- | --- | --- | --- |
| Experimental run 1 (1.1) | 4-7 July | 72 | 67 |
| Experimental run 1 (1.2) | 11-14 July | 71 | 39 |
| Experimental run 1 (1.3) | 14-21 July | 48 | 77 |
| Experimental run 1 (1.4) | 21-25 July | 55 | 51 |
| Experimental run 1 (1.5) | 25-28 July | 68 | 49 |
| Experimental run 1 (1.6) | 28-July-1 August | 69 | 60 |
| Experimental run 1 (1.7) | 1-11 August | 70 | 30 |
| Experimental run 2 (2.1) | 16-20 August | 72 | 71 |
| Experimental run 2 (2.2) | 24-28 August | 73 | 71 |
| Experimental run 2 (2.3) | 1-5 September | 72 | 72 |
| Experimental run 2 (2.4) | 9-13 September | 70 | 72 |
| Experimental run 2 (2.5) | 17-21 September | 71 | 72 |
| Experimental run 2 (2.6) | 25-29 September | 72 | 71 |
| Experimental run 2 (2.7) | 3-7 October | 71 | 72 |
| Experimental run 2 (2.8) | 7-13 October | 71 | 71 |

**Table S2.** Pairwise PERMANOVA results comparing differences in pollinator community composition among populations. Using an abundance matrix, we calculated Bray-Curtis distances to quantify the dissimilarities in pollinator assemblages. We performed pairwise tests between populations and adjusted P values to account for multiple comparisons.

| Site1 | Site2 | R<sup>2</sup> | P value |
| --- | --- | --- | --- |
| Even 1 | Clumped 1 | 0.12 | 0.364 |
| Even 1 | Clumped 2 | 0.72 | 0.002 |
| Even 1 | Even 2 | 0.66 | 0.002 |
| Clumped 1 | Clumped 2 | 0.77 | 0.001 |
| Clumped 1 | Even 2 | 0.67 | 0.004 |
| Clumped 2 | Even 2 | 0.09 | 0.307 |

**Table S3.**
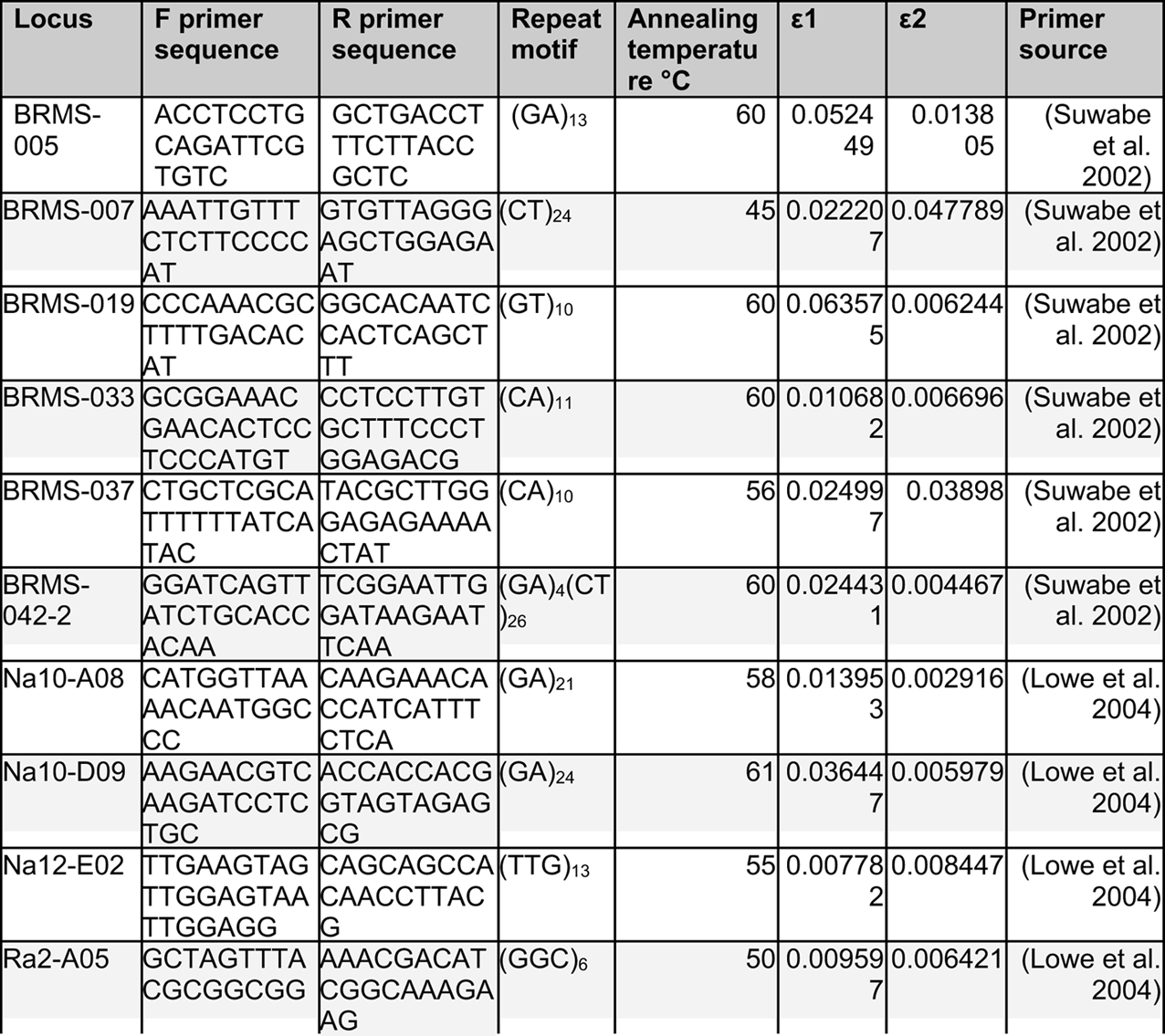
Primers used for microsatellite analysis. Information is provided on primer pairs used in this study including locus information, PCR parameters, and error rates.

**Table S4.**
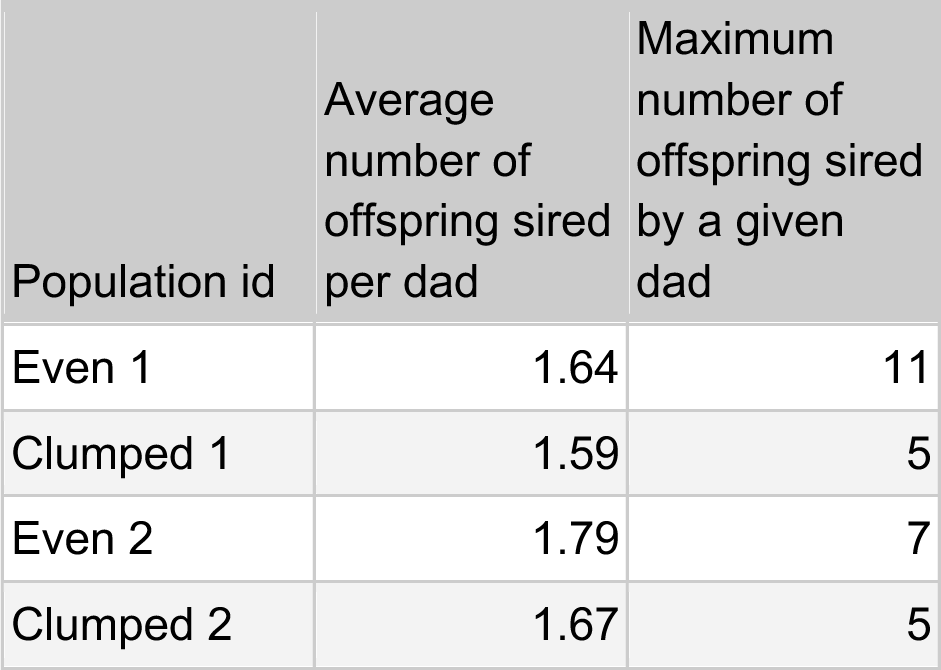
Paternity summaries per population including the average number of offspring sired per dad per plot and also the maximum number of offspring sired by any given dad within the plot across the entire season.

**Table S5.**
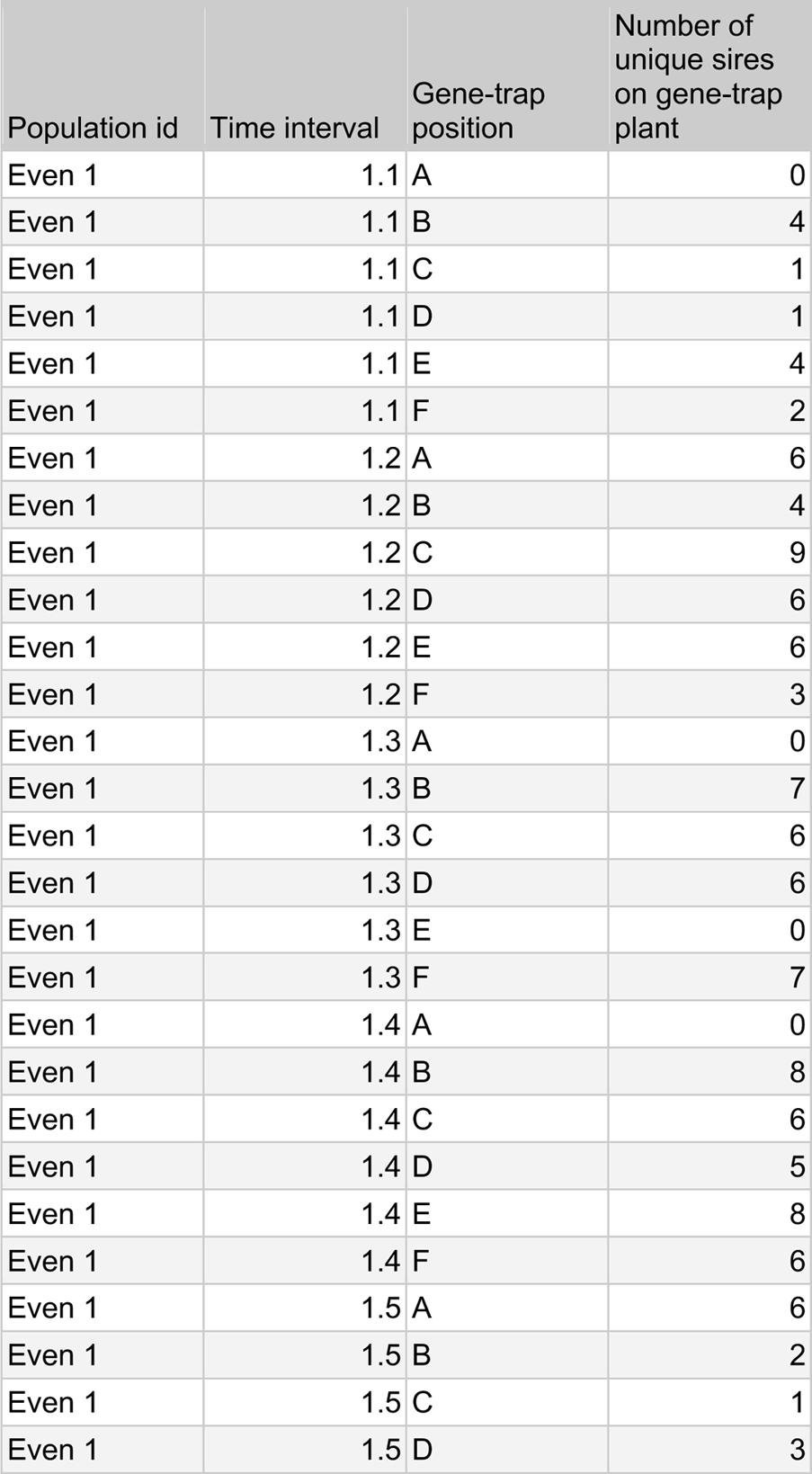
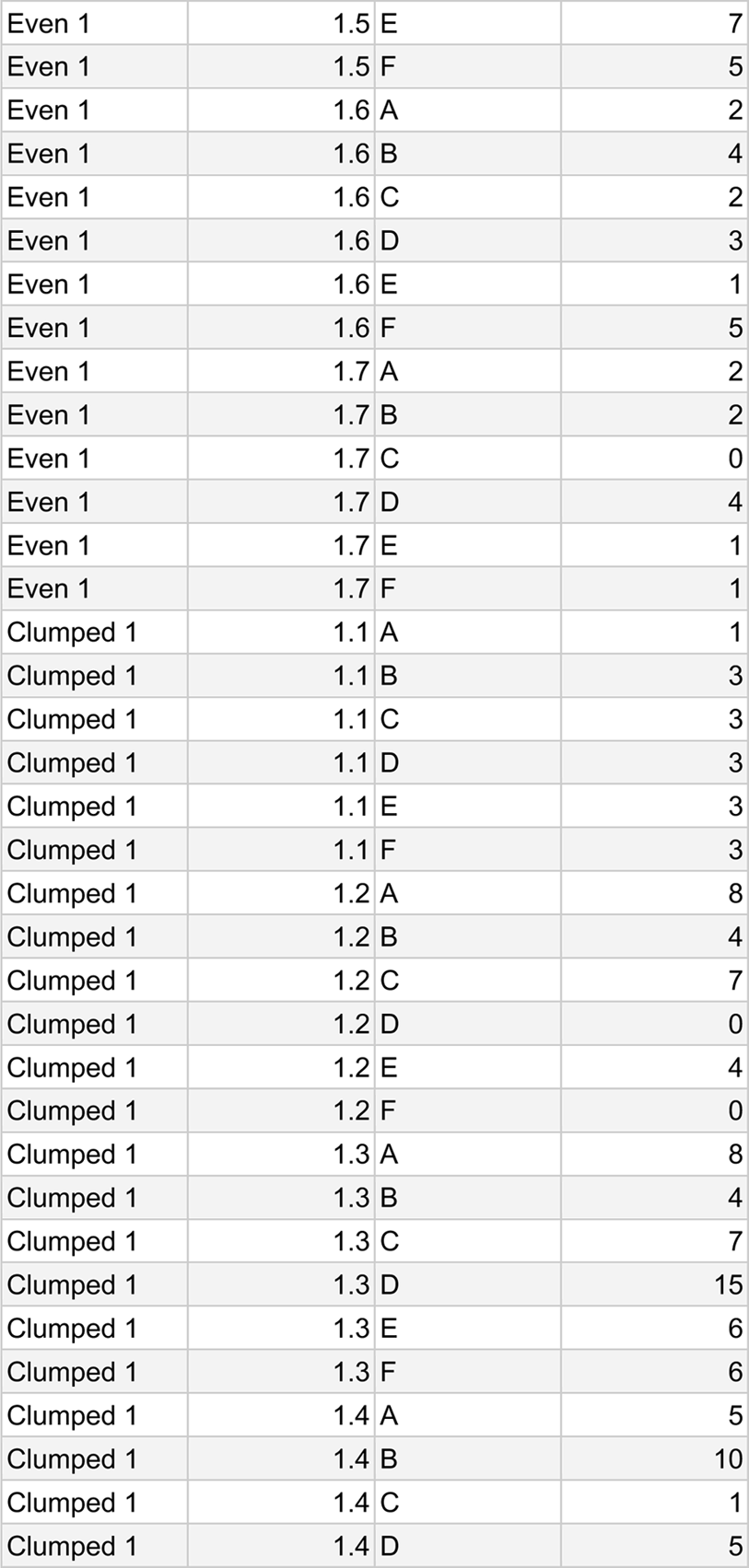
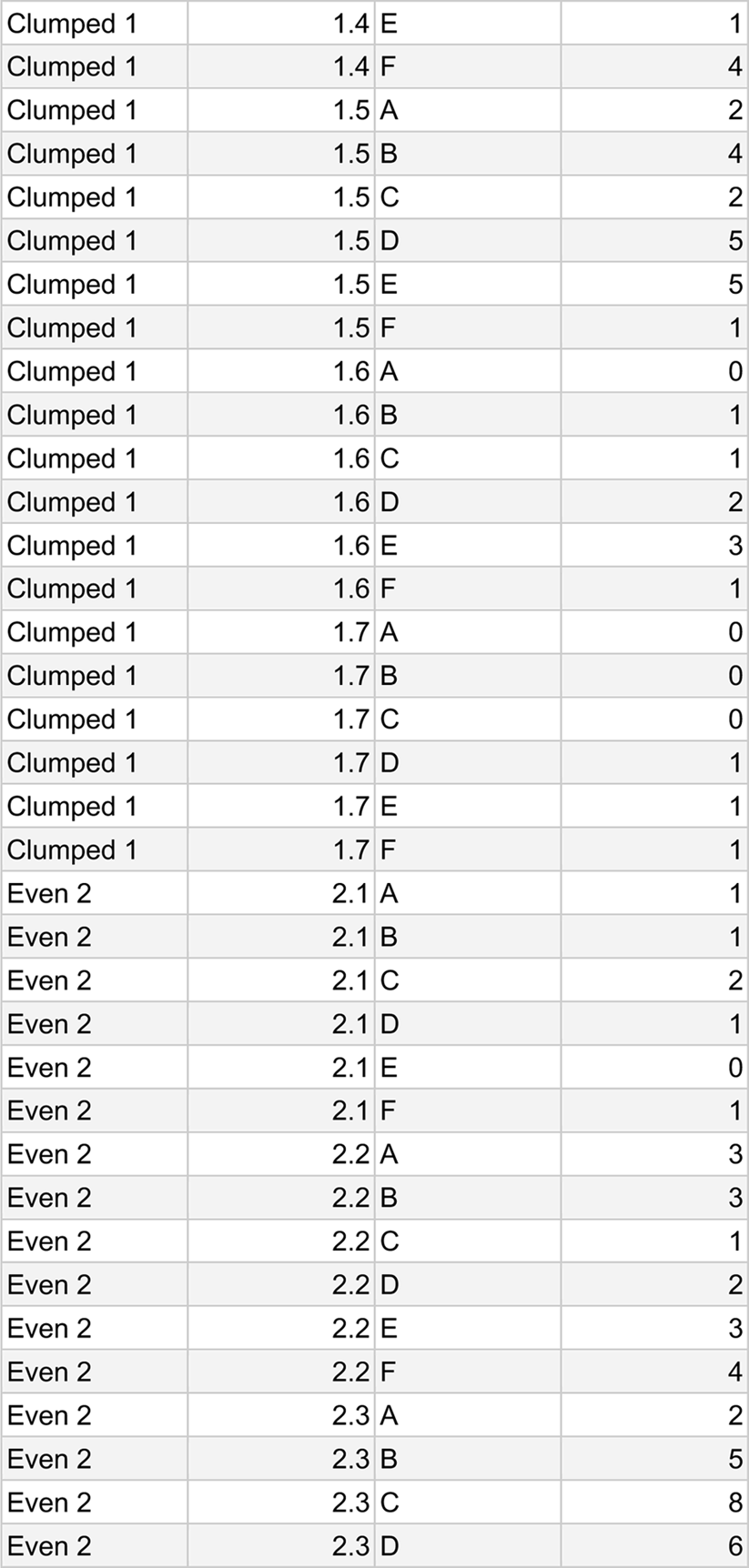
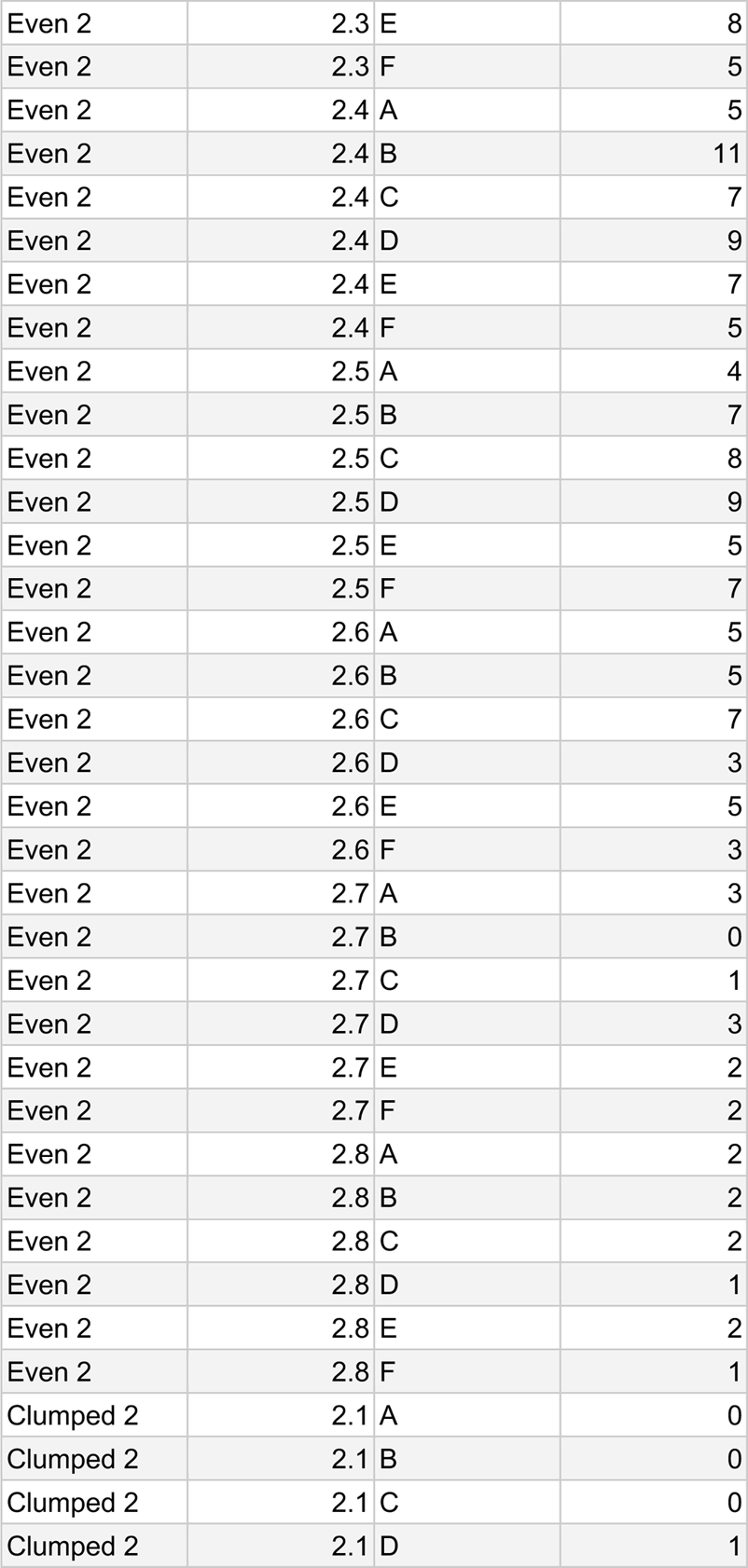
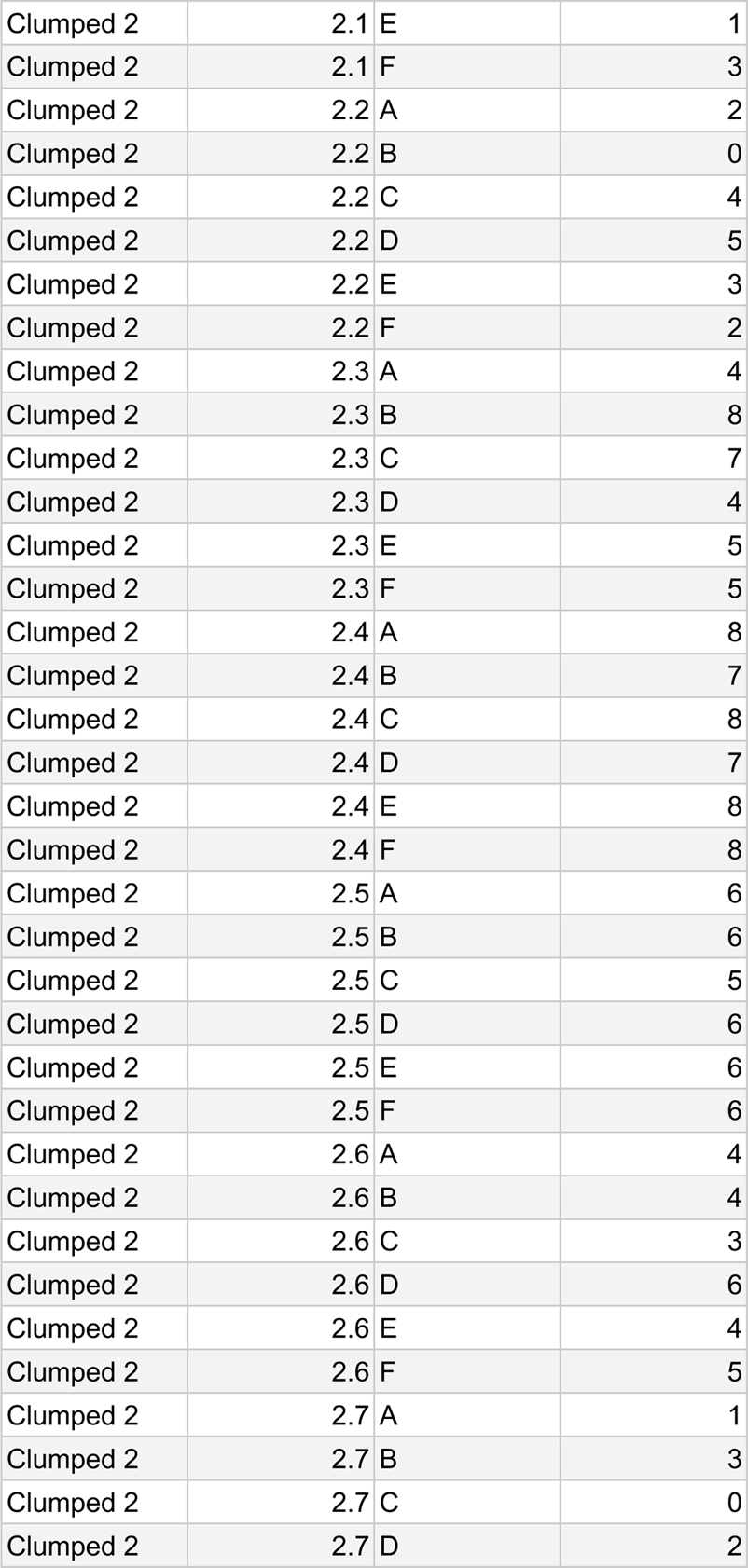
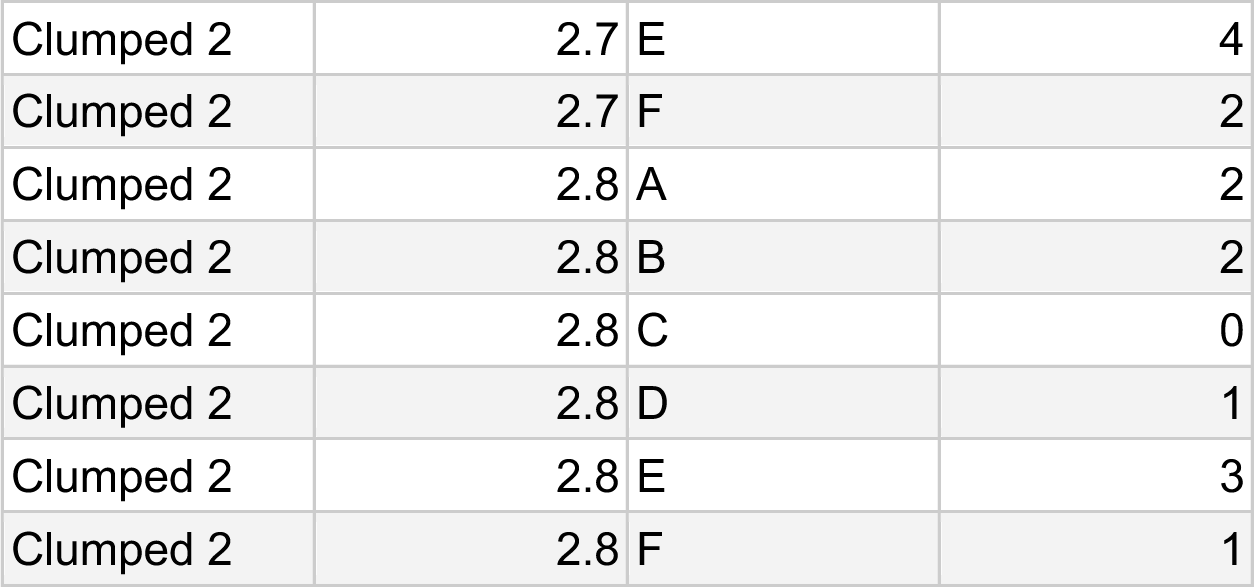
Diversity of mates on gene-trap plants. One gene-trap was added to each population at 6 positions over 7 intervals of time for experimental run 1 and 8 intervals of time for experimental run 2. The number of unique sires on gene-trap plants is a measure of the diversity of population sires that successfully fertilized an ovule on the gene-trap at that point in time.

**Table S6.** Longitudinal and cross-sectional estimates for selection across traits and plots (β [95% CI). gene-trap intervals can be related to dates in the season using Table S1. Clumped 2 interval 8 (gene-trap interval 2.8) was removed from analysis due to failing convergence and Gelman diagnostic.

| Phenotype | Selection gradient ($\beta$) [95% CI] | Cross-sectional or longitudinal selection gradient | Gene-trap interval | Population id |
| --- | --- | --- | --- | --- |
| flowering time | -0.239 [-0.400,-0.087] | cross sectional ($\beta$) | 1.1 | Even 1 |
| flowering time | -0.199 [-0.295,-0.114] | cross sectional ($\beta$) | 1.2 | Even 1 |
| flowering time | -0.055 [-0.143,0.026] | cross sectional ($\beta$) | 1.3 | Even 1 |
| flowering time | 0.005 [-0.072,0.079] | cross sectional ($\beta$) | 1.4 | Even 1 |
| flowering time | 0.020 [-0.043,0.080] | cross sectional ($\beta$) | 1.5 | Even 1 |
| flowering time | 0.081 [0.021,0.137] | cross sectional ($\beta$) | 1.6 | Even 1 |
| flowering time | 0.170 [0.104,0.239] | cross sectional ($\beta$) | 1.7 | Even 1 |
| flowering time | 0.010 [-0.030,0.043] | longitudinal ($\beta$) | n/a | Even 1 |
| flowering time | -0.376 [-0.576,-0.196] | cross sectional ($\beta$) | 1.1 | Clumped 1 |
| flowering time | 0.025 [-0.108,0.147] | cross sectional ($\beta$) | 1.2 | Clumped 1 |
| flowering time | -0.003 [-0.080,0.068] | cross sectional ($\beta$) | 1.3 | Clumped 1 |
| flowering time | -0.006 [-0.135,0.108] | cross sectional ($\beta$) | 1.4 | Clumped 1 |
| flowering time | 0.000 [-0.096,0.095] | cross sectional ($\beta$) | 1.5 | Clumped 1 |
| flowering time | 0.147 [0.054,0.253] | cross sectional ($\beta$) | 1.6 | Clumped 1 |
| flowering time | 0.081 [-0.102,0.321] | cross sectional ($\beta$) | 1.7 | Clumped 1 |
| flowering time | 0.038 [-0.010,0.080] | longitudinal ($\beta$) | n/a | Clumped 1 |
| flowering time | -24.367 [-34.563,-7.917] | cross sectional ($\beta$) | 2.1 | Even 2 |
| flowering time | -0.208 [-0.278,-0.136] | cross sectional ($\beta$) | 2.2 | Even 2 |
| flowering time | 0.014 [-0.040,0.067] | cross sectional ($\beta$) | 2.3 | Even 2 |
| flowering time | -0.003 [-0.056,0.051] | cross sectional ($\beta$) | 2.4 | Even 2 |
| flowering time | 0.079 [0.025,0.137] | cross sectional ($\beta$) | 2.5 | Even 2 |
| flowering time | 0.231 [0.149,0.319] | cross sectional ($\beta$) | 2.6 | Even 2 |
| flowering time | 0.086 [-0.010,0.213] | cross sectional ($\beta$) | 2.7 | Even 2 |
| flowering time | -0.065 [-0.174,0.064] | cross sectional ($\beta$) | 2.8 | Even 2 |
| flowering time | 0.040 [0.009,0.072] | longitudinal ($\beta$) | n/a | Even 2 |
| flowering time | -0.159 [-0.237,-0.082] | cross sectional ($\beta$) | 2.2 | Clumped 2 |
| flowering time | -0.048 [-0.096,0.000] | cross sectional ($\beta$) | 2.3 | Clumped 2 |
| flowering time | 0.047 [-0.004,0.099] | cross sectional ($\beta$) | 2.4 | Clumped 2 |
| flowering time | 0.104 [0.051,0.160] | cross sectional ($\beta$) | 2.5 | Clumped 2 |
| flowering time | 0.072 [0.024,0.122] | cross sectional ($\beta$) | 2.6 | Clumped 2 |
| flowering time | 0.180 [0.108,0.260] | cross sectional ($\beta$) | 2.7 | Clumped 2 |
| flowering time | 0.560 [0.381,0.775] | cross sectional ($\beta$) | 2.8 | Clumped 2 |
| flowering time | 0.022 [-0.005,0.049] | longitudinal ($\beta$) | n/a | Clumped 2 |
| duration | 0.051 [-0.027,0.117] | cross sectional ($\beta$) | 1.1 | Even 1 |
| duration | -0.019 [-0.088,0.044] | cross sectional ($\beta$) | 1.2 | Even 1 |
| duration | 0.025 [-0.044,0.083] | cross sectional ($\beta$) | 1.3 | Even 1 |
| duration | 0.009 [-0.054,0.066] | cross sectional ($\beta$) | 1.4 | Even 1 |
| duration | 0.041 [-0.020,0.097] | cross sectional ($\beta$) | 1.5 | Even 1 |
| duration | 0.093 [0.032,0.152] | cross sectional ($\beta$) | 1.6 | Even 1 |
| duration | 0.180 [0.115,0.248] | cross sectional ($\beta$) | 1.7 | Even 1 |
| duration | 0.005 [-0.026,0.034] | longitudinal ($\beta$) | n/a | Even 1 |
| duration | -0.280 [-0.431,-0.137] | cross sectional ($\beta$) | 1.1 | Clumped 1 |
| duration | 0.008 [-0.130,0.121] | cross sectional ($\beta$) | 1.2 | Clumped 1 |
| duration | -0.100 [-0.197,-0.009] | cross sectional ($\beta$) | 1.3 | Clumped 1 |
| duration | -0.035 [-0.134,0.053] | cross sectional ($\beta$) | 1.4 | Clumped 1 |
| duration | -0.004 [-0.095,0.075] | cross sectional ($\beta$) | 1.5 | Clumped 1 |
| duration | 0.154 [0.103,0.204] | cross sectional ($\beta$) | 1.6 | Clumped 1 |
| duration | 0.205 [0.084,0.372] | cross sectional ($\beta$) | 1.7 | Clumped 1 |
| duration | -0.025 [-0.075,0.022] | longitudinal ($\beta$) | n/a | Clumped 1 |
| duration | 0.045 [0.014,0.075] | cross sectional ($\beta$) | 2.1 | Even 2 |
| duration | 0.012 [-0.024,0.048] | cross sectional ($\beta$) | 2.2 | Even 2 |
| duration | 0.005 [-0.027,0.033] | cross sectional ($\beta$) | 2.3 | Even 2 |
| duration | 0.014 [-0.016,0.044] | cross sectional ($\beta$) | 2.4 | Even 2 |
| duration | -0.010 [-0.046,0.022] | cross sectional ($\beta$) | 2.5 | Even 2 |
| duration | -0.012 [-0.055,0.028] | cross sectional ($\beta$) | 2.6 | Even 2 |
| duration | 0.012 [-0.031,0.051] | cross sectional ($\beta$) | 2.7 | Even 2 |
| duration | 0.036 [-0.010,0.078] | cross sectional ($\beta$) | 2.8 | Even 2 |
| duration | 0.003 [-0.015,0.020] | longitudinal ($\beta$) | n/a | Even 2 |
| duration | 0.031 [-0.001,0.059] | cross sectional ($\beta$) | 2.2 | Clumped 2 |
| duration | -0.021 [-0.064,0.019] | cross sectional ($\beta$) | 2.3 | Clumped 2 |
| duration | 0.022 [-0.016,0.057] | cross sectional ($\beta$) | 2.4 | Clumped 2 |
| duration | 0.046 [0.011,0.082] | cross sectional ($\beta$) | 2.5 | Clumped 2 |
| duration | 0.109 [0.076,0.145] | cross sectional ($\beta$) | 2.6 | Clumped 2 |
| duration | 0.165 [0.097,0.233] | cross sectional ($\beta$) | 2.7 | Clumped 2 |
| duration | 0.484 [0.285,0.709] | cross sectional ($\beta$) | 2.8 | Clumped 2 |
| duration | 0.037 [0.020,0.055] | longitudinal ($\beta$) | n/a | Clumped 2 |
| total flowers | -0.001 [-0.013,0.010] | cross sectional ($\beta$) | 1.1 | Even 1 |
| total flowers | -0.003 [-0.015,0.007] | cross sectional ($\beta$) | 1.2 | Even 1 |
| total flowers | 0.005 [-0.007,0.017] | cross sectional ($\beta$) | 1.3 | Even 1 |
| total flowers | 0.012 [0.002,0.021] | cross sectional ($\beta$) | 1.4 | Even 1 |
| total flowers | 0.013 [0.006,0.019] | cross sectional ($\beta$) | 1.5 | Even 1 |
| total flowers | 0.014 [0.008,0.020] | cross sectional ($\beta$) | 1.6 | Even 1 |
| total flowers | 0.004 [-0.005,0.012] | cross sectional ($\beta$) | 1.7 | Even 1 |
| total flowers | 0.008 [0.003,0.012] | longitudinal ($\beta$) | n/a | Even 1 |
| total flowers | -0.033 [-0.066,0.001] | cross sectional ($\beta$) | 1.1 | Clumped 1 |
| total flowers | -0.012 [-0.043,0.012] | cross sectional ($\beta$) | 1.2 | Clumped 1 |
| total flowers | 0.005 [-0.014,0.023] | cross sectional ($\beta$) | 1.3 | Clumped 1 |
| total flowers | 0.034 [0.016,0.052] | cross sectional ($\beta$) | 1.4 | Clumped 1 |
| total flowers | 0.017 [0.000,0.035] | cross sectional ($\beta$) | 1.5 | Clumped 1 |
| total flowers | 0.002 [-0.014,0.021] | cross sectional ($\beta$) | 1.6 | Clumped 1 |
| total flowers | 0.025 [-0.003,0.064] | cross sectional ($\beta$) | 1.7 | Clumped 1 |
| total flowers | 0.007 [-0.004,0.017] | longitudinal ($\beta$) | n/a | Clumped 1 |
| total flowers | 0.015 [-0.004,0.034] | cross sectional ($\beta$) | 2.1 | Even 2 |
| total flowers | 0.018 [0.002,0.033] | cross sectional ($\beta$) | 2.2 | Even 2 |
| total flowers | -0.001 [-0.012,0.011] | cross sectional ($\beta$) | 2.3 | Even 2 |
| total flowers | 0.015 [0.004,0.024] | cross sectional ($\beta$) | 2.4 | Even 2 |
| total flowers | 0.015 [0.005,0.024] | cross sectional ($\beta$) | 2.5 | Even 2 |
| total flowers | 0.012 [0.001,0.021] | cross sectional ($\beta$) | 2.6 | Even 2 |
| total flowers | 0.014 [0.006,0.022] | cross sectional ($\beta$) | 2.7 | Even 2 |
| total flowers | 0.023 [0.015,0.029] | cross sectional ($\beta$) | 2.8 | Even 2 |
| total flowers | 0.007 [0.001,0.013] | longitudinal ($\beta$) | n/a | Even 2 |
| total flowers | 0.005 [-0.005,0.014] | cross sectional ($\beta$) | 2.2 | Clumped 2 |
| total flowers | 0.009 [-0.002,0.020] | cross sectional ($\beta$) | 2.3 | Clumped 2 |
| total flowers | 0.006 [-0.006,0.016] | cross sectional ($\beta$) | 2.4 | Clumped 2 |
| total flowers | 0.020 [0.011,0.030] | cross sectional ($\beta$) | 2.5 | Clumped 2 |
| total flowers | 0.012 [0.003,0.022] | cross sectional ($\beta$) | 2.6 | Clumped 2 |
| total flowers | -0.001 [-0.011,0.009] | cross sectional ($\beta$) | 2.7 | Clumped 2 |
| total flowers | 0.012 [0.003,0.021] | cross sectional ($\beta$) | 2.8 | Clumped 2 |
| total flowers | 0.007 [0.001,0.013] | longitudinal ($\beta$) | n/a | Clumped 2 |

**Table S7.** Distance decay coefficients [95% CI] across populations and intervals of time. Clumped 2 interval 8 (gene-trap interval 2.8) was removed from analysis due to failing convergence and Gelman diagnostic.

| Coefficient of distance decay<br>[95% CI] | Population id | gene-trap interval |
| --- | --- | --- |
| -0.54 [-0.81,-0.29] | Even 1 | 1.1 |
| -1.74 [-2.21,-1.31] | Even 1 | 1.2 |
| -1.16 [-1.69,-0.72] | Even 1 | 1.3 |
| -0.98 [-1.34,-0.68] | Even 1 | 1.4 |
| -0.55 [-0.80,-0.33] | Even 1 | 1.5 |
| -0.52 [-0.77,-0.30] | Even 1 | 1.6 |
| -0.51 [-0.76,-0.28] | Even 1 | 1.7 |
| -1.19 [-1.38,-1.00] | Even 1 | n/a |
| -0.51 [-0.67,-0.37] | Clumped 1 | 1.1 |
| -0.32 [-0.53,-0.14] | Clumped 1 | 1.2 |
| -0.27 [-0.39,-0.14] | Clumped 1 | 1.3 |
| -0.51 [-0.75,-0.31] | Clumped 1 | 1.4 |
| -0.27 [-0.41,-0.12] | Clumped 1 | 1.5 |
| -0.10 [-0.24,0.05] | Clumped 1 | 1.6 |
| -0.43 [-0.66,-0.23] | Clumped 1 | 1.7 |
| -0.30 [-0.38,-0.23] | Clumped 1 | n/a |
| -0.42 [-0.61,-0.22] | Even 2 | 2.1 |
| -0.35 [-0.58,-0.14] | Even 2 | 2.2 |
| -1.11 [-1.48,-0.78] | Even 2 | 2.3 |
| -0.65 [-0.88,-0.43] | Even 2 | 2.4 |
| -0.39 [-0.62,-0.19] | Even 2 | 2.5 |
| -0.61 [-0.86,-0.37] | Even 2 | 2.6 |
| -0.46 [-0.68,-0.24] | Even 2 | 2.7 |
| -0.22 [-0.44,-0.01] | Even 2 | 2.8 |
| -0.59 [-0.72,-0.47] | Even 2 | n/a |
| -0.40 [-0.54,-0.26] | Clumped 2 | 2.2 |
| -0.63 [-0.84,-0.47] | Clumped 2 | 2.3 |
| -0.45 [-0.60,-0.32] | Clumped 2 | 2.4 |
| -0.33 [-0.45,-0.22] | Clumped 2 | 2.5 |
| -0.35 [-0.46,-0.23] | Clumped 2 | 2.6 |
| -0.37 [-0.50,-0.25] | Clumped 2 | 2.7 |
| -0.44 [-0.57,-0.30] | Clumped 2 | 2.8 |
| -0.42 [-0.50,-0.35] | Clumped 2 | n/a |

